# MEF2C controls segment-specific gene regulatory networks that direct heart tube morphogenesis

**DOI:** 10.1101/2024.11.01.621613

**Authors:** Jonathon M. Muncie-Vasic, Tanvi Sinha, Alexander P. Clark, Emily F. Brower, Jeffrey J. Saucerman, Brian L. Black, Benoit G. Bruneau

## Abstract

The gene regulatory networks (GRNs) that control early heart formation are beginning to be understood, but lineage-specific GRNs remain largely undefined. We investigated networks controlled by the vital transcription factor MEF2C, with a time course of single-nucleus RNA- and ATAC-sequencing in wild-type and *Mef2c*-null embryos. We identified a “posteriorized” cardiac gene signature and chromatin landscape in the absence of MEF2C. Integrating our multiomics data in a deep learning-based model, we constructed developmental trajectories for each of the outflow tract, ventricular, and inflow tract segments, and alterations of these in *Mef2c*-null embryos. We computationally identified segment-specific MEF2C-dependent enhancers, with activity in the developing zebrafish heart. Finally, using inferred GRNs we discovered that the *Mef2c*-null heart malformations are partly driven by increased activity of the nuclear hormone receptor NR2F2. Our results delineate lineage-specific GRNs in the early heart tube and provide a generalizable framework for dissecting transcriptional networks governing developmental processes.

---

A recurrent observation in developmental biology is that the same transcription factor (TF) can play different roles depending on the cells or developmental stage in which it is expressed. The cell type- or stage-specific activity of a TF is dependent on a number of factors, including its own expression level, the availability of potential co-factors, the presence or absence of other TFs that may act co-operatively or antagonistically, and the dynamic state of the chromatin landscape in which the TF is acting^1–4^. Together, these phenomena can be conceptualized as transcriptional or gene regulatory networks (GRNs) that can be used to understand the role of a particular TF within a given developmental context^5^. However, for most TFs, the precise GRNs underlying their pleiotropic activity in distinct cell types and at different developmental timepoints remain undefined.

The developing heart presents an excellent model for exploring this question. Several TFs critical for cardiac development have been identified and their expression patterns are well-defined^6–8^. Moreover, lineage tracing studies have revealed distinct progenitor cell types that contribute to the various segments of the linear heart tube^7–11^ (Fig. 1a). To explore in detail how a specific TF can have distinct roles within different parts of a developing organ, we focused on a key cardiac transcription factor, MEF2C^12^, and its role in each of the developing heart tube segments. In mice, loss of MEF2C leads to cardiac defects and embryonic lethality by mid-gestation at embryonic day (E) 10.5^13^. Interestingly, although MEF2C is expressed in both first heart field (FHF) and second heart field (SHF) progenitors in the cardiac crescent at E7.75, and throughout the developing heart tube from E8.5 to E9, the loss of MEF2C causes distinct defects in different segments of the heart tube (Fig. 1b-c, Extended Data Fig. 1a). At E9, the outflow tract, which gives rise to the aorta and pulmonary trunk, is severely hypoplastic and only a single hypoplastic ventricle has formed. By contrast, the inflow tract, precursor to the atria, appears to be expanded and improperly patterned (Fig. 1c). These segment-specific defects suggest that MEF2C plays distinct regulatory roles in the outflow tract, ventricles, and inflow tract.

**Figure 1:**
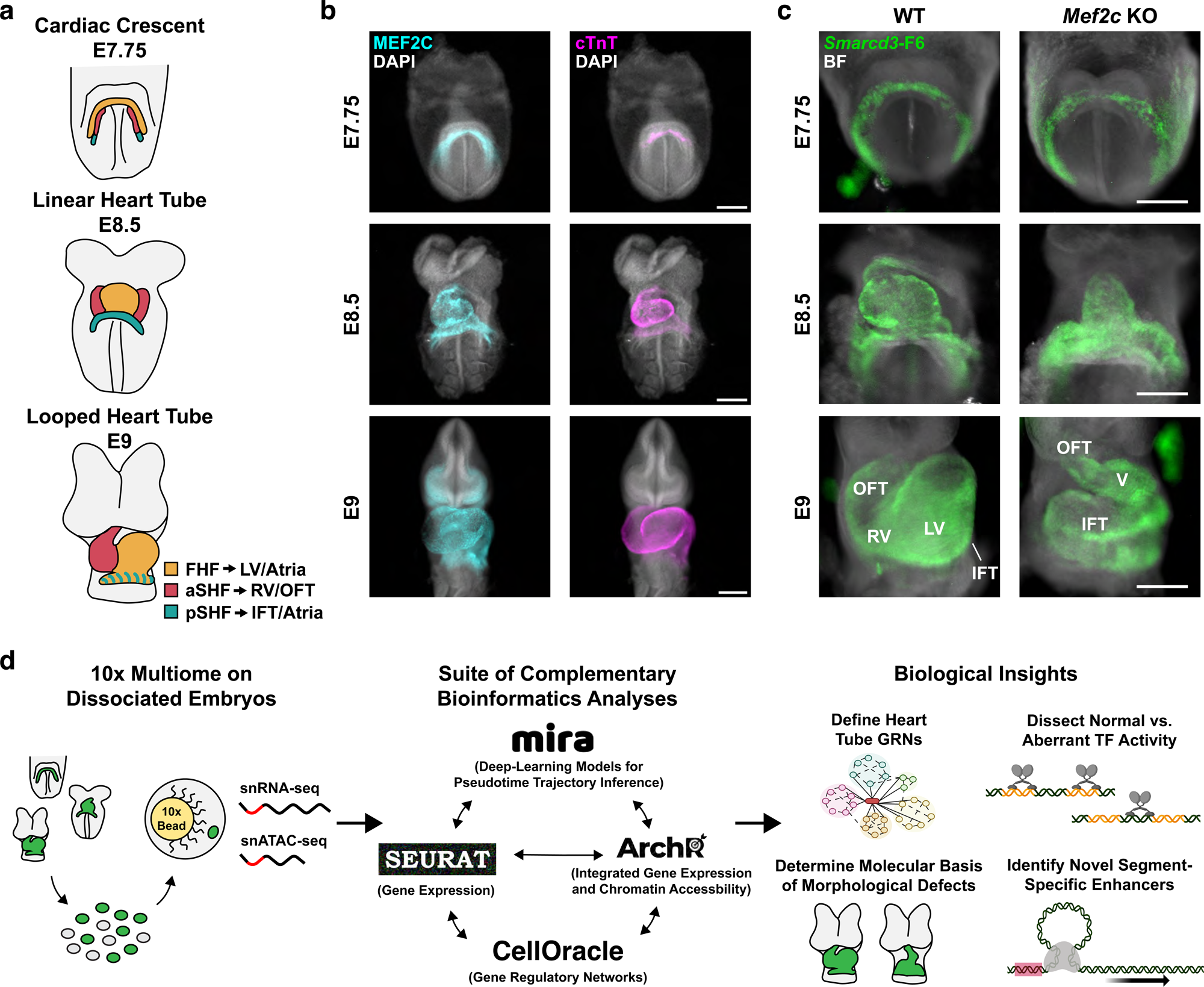
MEF2C is expressed throughout the developing heart tube and its loss causes segment-specific defects. a) Schematic of cardiac progenitors and their contributions to linear heart tube development from cardiac crescent (E7.75) to looped heart tube (E9) stage. b) Immunofluorescent staining of MEF2C (cyan) and cardiac Troponin T (cTnT, magenta) in E7.75, E8.5, and E9 WT embryos. c) Representative images of WT and *Mef2c* KO embryos at E7.75, E8.5 and E9. Cardiac progenitors are marked by the *Smarcd3*-F6-eGFP reporter transgene (green). BF, brightfield. d) Schematic of the methodology and biological insights presented in the current study. Elements of this panel were created in BioRender. B, B. (2024) https://BioRender.com/i72e213. Scale bars = 200 μm. FHF, first heart field; aSHF, anterior second heart field; pSHF, posterior second heart field; LV, left ventricle; RV, right ventricle; V, ventricle; IFT, inflow tract; OFT, outflow tract.

To dissect the segment-specific regulation of heart tube morphogenesis by MEF2C, we performed combined single-nucleus RNA-seq and ATAC-seq (snRNA-seq and snATAC-seq) on wild-type (WT) and *Mef2c* knockout (KO) mouse embryos at key stages throughout heart tube development. Taking advantage of the ability to match the transcriptomic and chromatin accessibility data for every cell in the dataset, we applied a suite of complementary bioinformatics tools to construct GRNs for cardiac progenitor lineages, precisely defined the role of MEF2C in regulating gene expression and chromatin accessibility in the developing heart tube, identified novel MEF2C-dependent enhancers, and discovered a genetic interaction that underlies the inflow tract malformation in *Mef2c* KO embryos (Fig. 1d).

## RESULTS

### Loss of MEF2C reduces expression of key cardiomyocyte genes across all heart tube segments and alters expression of anterior/posterior markers

To assess the role of MEF2C in controlling gene expression in the developing heart tube, we performed snRNA-seq on stage-matched WT and *Mef2c* KO embryos at E7.75, E8.5, and E9 (n = 2 embryos for each genotype at each stage, Extended Data Fig. 1b). All embryos carried the *Smarcd3*-F6-eGFP transgene reporter to mark cardiac progenitors^14^. At E7.75, we harvested the entire embryos, whereas at E8.5 and E9 we removed the headfolds and posterior trunk via microdissection to increase the proportion of cardiac progenitors captured for sequencing. We created separate Seurat objects^15^ for the embryos profiled at each timepoint (E7.75, E8.5, and E9). Following pre-processing, quality control, normalization with scTransform^16^, and dimensionality reduction, we performed unbiased Louvain clustering and plotted the data in uniform manifold approximation and projection (UMAP) space. We then used the expression of known marker genes to identify the cell types represented by each cluster (Extended Data Fig. 2a-f, Supplementary Table 1).

To resolve specific cardiomyocyte subtypes, we subset and re-clustered cell types of interest, including cardiac progenitors, cardiomyocytes, and related mesoderm cell types (Fig. 2a-c, Extended Data Fig. 2g-i, Supplementary Table 2). By doing so, we were able to identify clusters representing FHF and early differentiating cardiomyocytes (CMs/FHF), SHF, and juxtacardiac field (JCF) progenitors^17^ at E7.75 (Fig. 2a). At E8.5, we could distinguish inflow tract cardiomyocytes (IFT-CMs), ventricular cardiomyocytes (V-CMs), and outflow tract cardiomyocytes (OFT-CMs) (Fig. 2b). At E9 we identified the IFT-CMs more specifically as either atrial or atrioventricular canal (A- or AVC-CMs) (Fig. 2c). We next performed differential gene expression analysis between *Mef2c* KO and WT cells within each of these cell types (Fig. 2d-f, Supplementary Table 3). Overall, more genes were down-regulated than up-regulated in the *Mef2c* KO relative to WT embryos, consistent with the notion that MEF2C is an activator of the myogenic transcriptional program^18^. Among the top differentially expressed genes (DEGs), we found several genes that have been associated with high confidence to congenital heart defects^19^, including *Myh6*, *Myh7*, *Actc1*, and *Foxp1* (Extended Data Fig. 2j). Furthermore, nearly all these top DEGs were identified as the nearest genes to MEF2C-occupied peaks in a previously published MEF2C ChIP-seq dataset of embryonic hearts at E14^20^, suggesting that these genes are direct targets of MEF2C in the developing heart.

**Figure 2:**
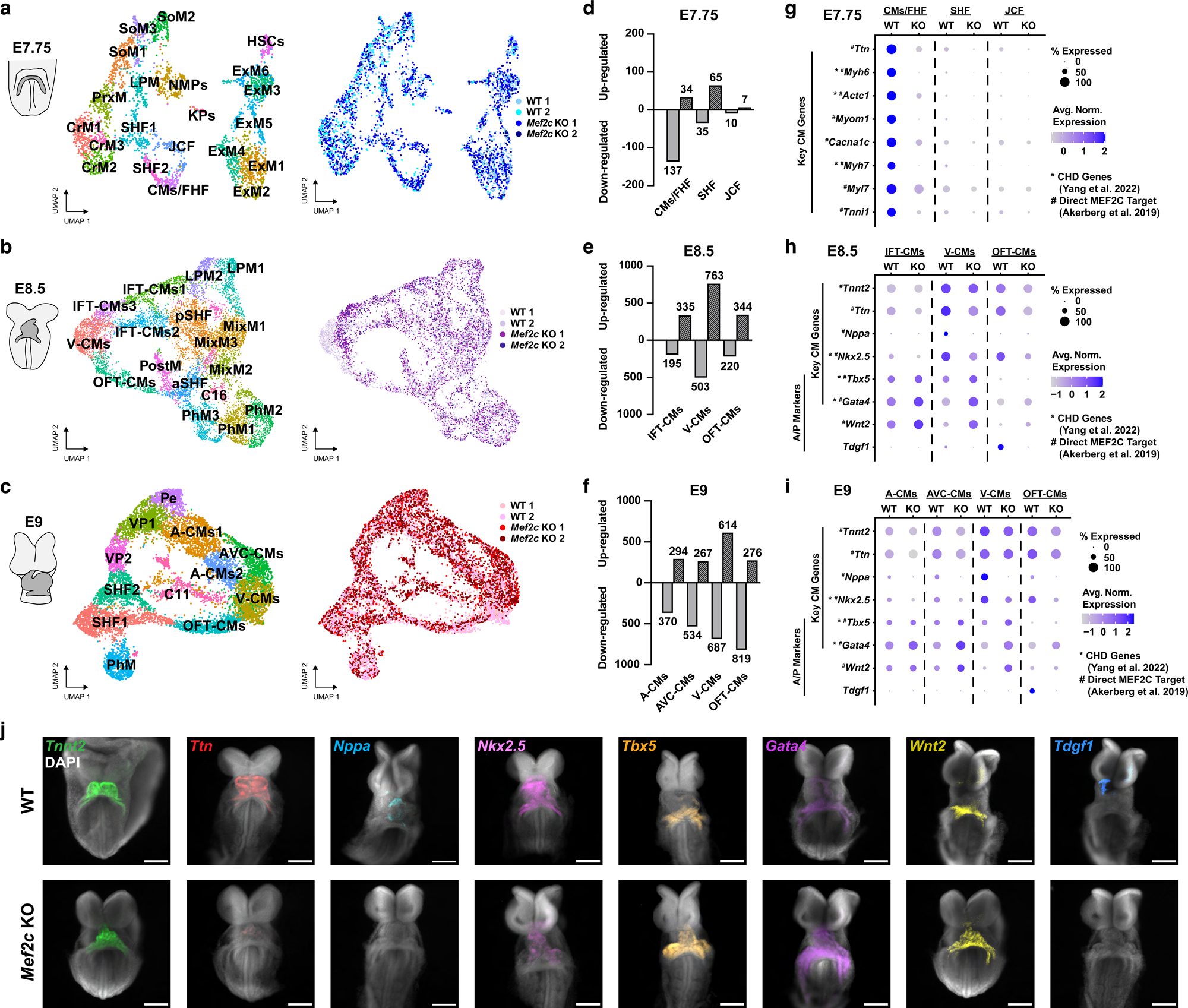
Loss of MEF2C reduces expression of key cardiomyocyte genes across all heart tube segments and alters expression of anterior/posterior markers. a-c) UMAPs of snRNA-seq data for cardiac progenitors, cardiomyocytes, and related mesoderm subtypes from E7.75 (a), E8.5 (b), and E9 (c) embryos labeled by cell type (left) and genotype/sample ID (right). d-f) Bar plots displaying the number of up-regulated and down-regulated genes in *Mef2c* KO relative to WT in cell types of interest at E7.75 (d), E8.5 (e), and E9 (f). g-i) Dot plots displaying expression of key CM genes and anterior/posterior (A/P) markers at E7.75 (g), E8.5 (h), and E9 (i). j) Fluorescence *in situ* hybridization of key CM genes and A/P markers in E8.5-E9 WT and *Mef2c* KO embryos. Note the reduced expression of CM genes *Tnnt2*, *Ttn*, *Nppa*, and *Nkx2-5*, the expanded expression of posterior markers *Tbx5*, *Gata4*, and *Wnt2*, and the loss of anterior OFT marker *Tdgf1* in *Mef2c* KO embryos compared to WT. Scale bars = 200 μm. CMs, cardiomyocytes; FHF, first heart field; SHF, second heart field; JCF, juxtacardiac field; CrM, cranial mesoderm; PrxM, paraxial mesoderm; LPM, lateral plate mesoderm; SoM, somitic mesoderm; NMPs, neuromesodermal progenitors; KPs, kidney progenitors; ExM, extraembryonic mesoderm; HSCs, hematopoietic stem cells; V-CMs, ventricular cardiomyocytes; IFT-CMs, inflow tract cardiomyocytes; OFT-CMs, outflow tract cardiomyocytes; aSHF, anterior second heart field; pSHF, posterior second heart field; PostM, posterior mesoderm; PhM, pharyngeal mesoderm; MixM, mixed mesoderm; A-CMs, atrial cardiomyocytes; AVC-CMs, atrioventricular canal cardiomyocytes; Pe, proepicardium; VP, venous pole; *, Genes known to be associated with CHDs^19^; #, Direct targets of MEF2C based on MEF2C ChIP-seq data^20^.

Closer examination of our DEG data (Supplementary Table 3) revealed two categories of DEGs. The first category consisted of key cardiomyocyte genes that were substantially down-regulated across all three heart tube segments in the *Mef2c* KO embryos, including genes encoding sarcomeric proteins such as *Tnnt2* and *Ttn,* the cardiac transcription factor *Nkx2-5*, and the natriuretic peptide *Nppa* (Fig. 2g-i). These data confirm the importance of MEF2C in early cardiomyocyte fate specification, as a direct activator of contractile protein gene expression and by regulating additional TFs and functional proteins necessary for proper cardiomyocyte development. The second category of DEGs consisted of genes involved in anterior/posterior patterning of the heart tube. Specifically, genes such as *Tbx5*, *Gata4*, and *Wnt2,* which are normally expressed only in the posterior IFT segment of the heart tube, were not only up-regulated in the IFT-CMs, but their expression was also expanded into the V-CMs in *Mef2c* KO (Fig. 2g-i). Additionally, expression of the anterior OFT-specific gene *Tdgf1* was completely lost in *Mef2c* KO (Fig. 2h-i), as previously demonstrated^21^. Consistent with this apparent posteriorization of gene expression in the heart tube, we also observed a larger proportion of A-CMs and AVC-CMs in *Mef2c* KO embryos compared to WT at E9, with a concomitant reduction in the proportion of V-CMs and OFT-CMs (Extended Data Fig. 2k). We validated the observed changes in gene expression with whole-mount embryo fluorescence *in situ* hybridization, confirming that *Mef2c* KO leads to broad reduction in the cardiomyocyte transcriptional program and a notable posteriorization of the developing heart tube (Fig. 2j).

### MEF2C regulates chromatin accessibility broadly throughout the heart tube and in a segment-specific manner

To investigate how MEF2C regulates the observed segment-specific gene expression changes, we integrated our chromatin accessibility and gene expression datasets using ArchR^22^. We performed dimensionality reduction using ArchR’s latent semantic index (LSI) implementation on the snRNA-seq and snATAC-seq modalities and then plotted the cells in UMAP space using the combined dimensions (Extended Data Fig. 3a-c). Matching cell barcodes in the snRNA-seq and snATAC-seq libraries allowed us to transfer the cell type labels from the Seurat objects to the integrated ArchR datasets. As we did with the gene expression data, we subset the cardiac cell types and associated mesoderm in order to better resolve cardiomyocyte subtypes. Finally, we created pseudobulk groups consisting of WT and *Mef2c* KO cells for each given cell type label in the subset datasets and called peaks representing accessible regions using MACS2^23^.

Our initial analyses revealed few changes to chromatin accessibility at E7.75 (Extended Data Fig. 3d). This was not entirely unexpected, given that there is no discernable phenotype in the *Mef2c* KO embryos at this timepoint, and agrees with the relatively few changes in gene expression we observed in CMs/FHF, SHF, and JCF cells at E7.75 (Fig. 2d). Thus, we focused on the E8.5 dataset to identify the earliest substantial changes in chromatin accessibility induced by the loss of MEF2C (Fig. 3a). Using our pseudobulked data, we identified differentially accessible regions (DARs) of chromatin between *Mef2c* KO and WT cells in each of the subset cell types and found that, as expected, cardiomyocytes displayed the largest numbers of DARs (Fig. 3b). We also found that in each of the CM subtypes (IFT-CMs, V-CMs, and OFT-CMs), *Mef2c* KO CMs lost more DARs than they gained (Fig. 3b), and these lost DARs were highly enriched for MEF2 binding motifs (Extended Data Fig. 3e). These data further reinforce MEF2C’s role as an activator of the contractile gene transcriptional program. Interestingly, when we examined the lost DARs in *Mef2c* KO embryos, we found 675 regions that demonstrated lost accessibility in two or more heart tube segments, and 2,549 regions that lost accessibility in a segment-specific manner (Fig. 3c). The former are likely regions involved in the regulation of general cardiomyocyte genes, while the latter may control segment-specific gene regulation. By contrast, the gained DARs in *Mef2c* KO embryos were almost exclusively unique to each heart tube segment (Fig. 3c). These segment-specific DARs are interesting because they provide evidence that distinct transcriptional networks exist for each of the three heart tube segments and that each segment-specific network is uniquely altered by the loss of MEF2C.

**Figure 3:**
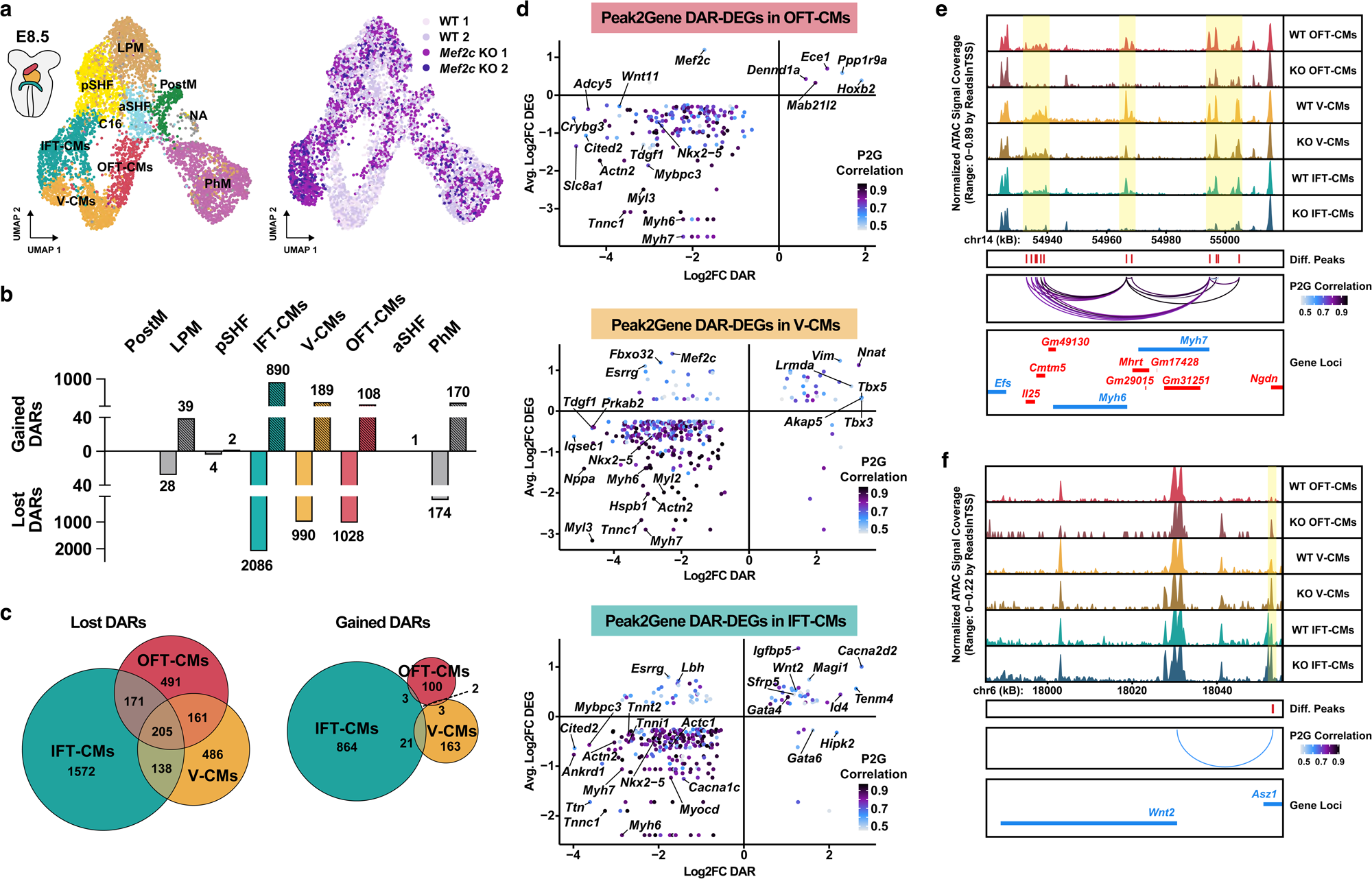
MEF2C regulates chromatin accessibility broadly throughout the heart tube and in a segment-specific manner. a) UMAP of integrated snRNA-seq and snATAC-seq data for cardiac progenitors, cardiomyocytes, and related mesoderm subtypes at E8.5 labeled by cell types determined from snRNA-seq clustering (left) and genotype/sample ID (right). b) Bar plots displaying the number of gained and lost DARs in *Mef2c* KO relative to WT cell types of interest at E8.5. c) Venn diagrams displaying unique and overlapping DARs in the three heart tube segments. d) Scatter plots displaying the relationship between the Log2 fold change (Log2FC) values of the DEG and DAR analyses for all identified Peak2Gene (P2G) links in the three heart tube segments. Dots are colored by the P2G correlation score. e-f) Genome browser tracks displaying snATAC-seq accessibility profiles at the *Myh6*/*Myh7* (e) and *Wnt2* (f) loci for the indicated pseudobulked cell types. DARs are highlighted and the peaks are indicated by red bars (*Mef2c* KO-vs-WT IFT-CMs). Loops indicate P2G links, colored by the P2G correlation score. V-CMs, ventricular cardiomyocytes; IFT-CMs, inflow tract cardiomyocytes; OFT-CMs, outflow tract cardiomyocytes; aSHF, anterior second heart field; pSHF, posterior second heart field; LPM, lateral plate mesoderm; PostM, posterior mesoderm; PhM, pharyngeal mesoderm; NA, cells not available in the snRNA-seq dataset; DAR, differentially accessible region; DEG, differentially expressed gene.

To identify the DARs most likely to represent *bona fide* regulatory elements, we performed a peak-to-gene linkage (P2G) analysis, which identifies statistically significant correlations between the accessibility of individual peaks and the expression of potential target genes within 250 kilobases of each peak. We found 817 P2G links in the OFT-CMs, 1,201 in the V-CMs, and 2,257 in the IFT-CMs. We then intersected the P2G links with the MEF2C-dependent DEGs at each timepoint (Supplementary Table 4) and plotted each P2G link according to the fold change of the peak’s accessibility, the fold change of linked gene’s differential expression, and the correlation strength of the P2G link (Fig. 3d). These P2G DAR-DEG correlation plots reveal a number of interesting observations. Genes such as *Nkx2-5*, *Myh6*, and *Myh7*, whose expression decreased throughout the heart tube in *Mef2c* KO embryos, are linked to DARs in all three segments (Fig. 3d). In particular, the *Myh6*/*Myh7* locus contains multiple regions that have P2G links and clearly altered accessibility in OFT-CMs, V-CMs, and IFT-CMs (Fig. 3e). This suggests that these DARs are likely to be regulatory elements that control expression of *Myh6* and *Myh7* and are sensitive to the expression of *Mef2c* in each segment of the heart tube. By contrast, there are also P2G links between peaks with lost accessibility and decreased gene expression that are unique to the individual segments, such as links to *Wnt11* in the OFT-CMs, *Nppa* in the V-CMs, and *Cacna1c* in the IFT-CMs (Fig. 3d). Furthermore, examining the links between peaks with gained accessibility and genes with increased expression in *Mef2c* KO reveals additional evidence for the posteriorized transcriptional program we previously described. Namely, P2G links to *Tbx5* in the V-CMs, as well as *Gata4* and *Wnt2* in the IFT-CMs (Fig. 3d and 3f). These P2G links represent potential regulatory elements that are activated or over-activated in response to the loss of MEF2C and, at least in part, promote the increased posteriorization of the heart tube.

### Each heart tube segment exhibits a distinct MEF2C-depedent developmental trajectory

Our multiomic datasets containing precisely staged embryos at different developmental timepoints provided us the opportunity to dissect how lineage trajectories unfold in WT embryos and are altered by loss of MEF2C. To this end, we applied multimodal Models for Integrated Regulatory Analysis (MIRA)^24^, which uses deep learning and probabilistic graphical models to construct sets of “topics” that represent either gene expression or peak accessibility. These topics can then be used to plot cells in UMAP space, assign cell identities, perform pseudotime analyses, and construct cell state trees (Fig. 4a).

**Figure 4:**
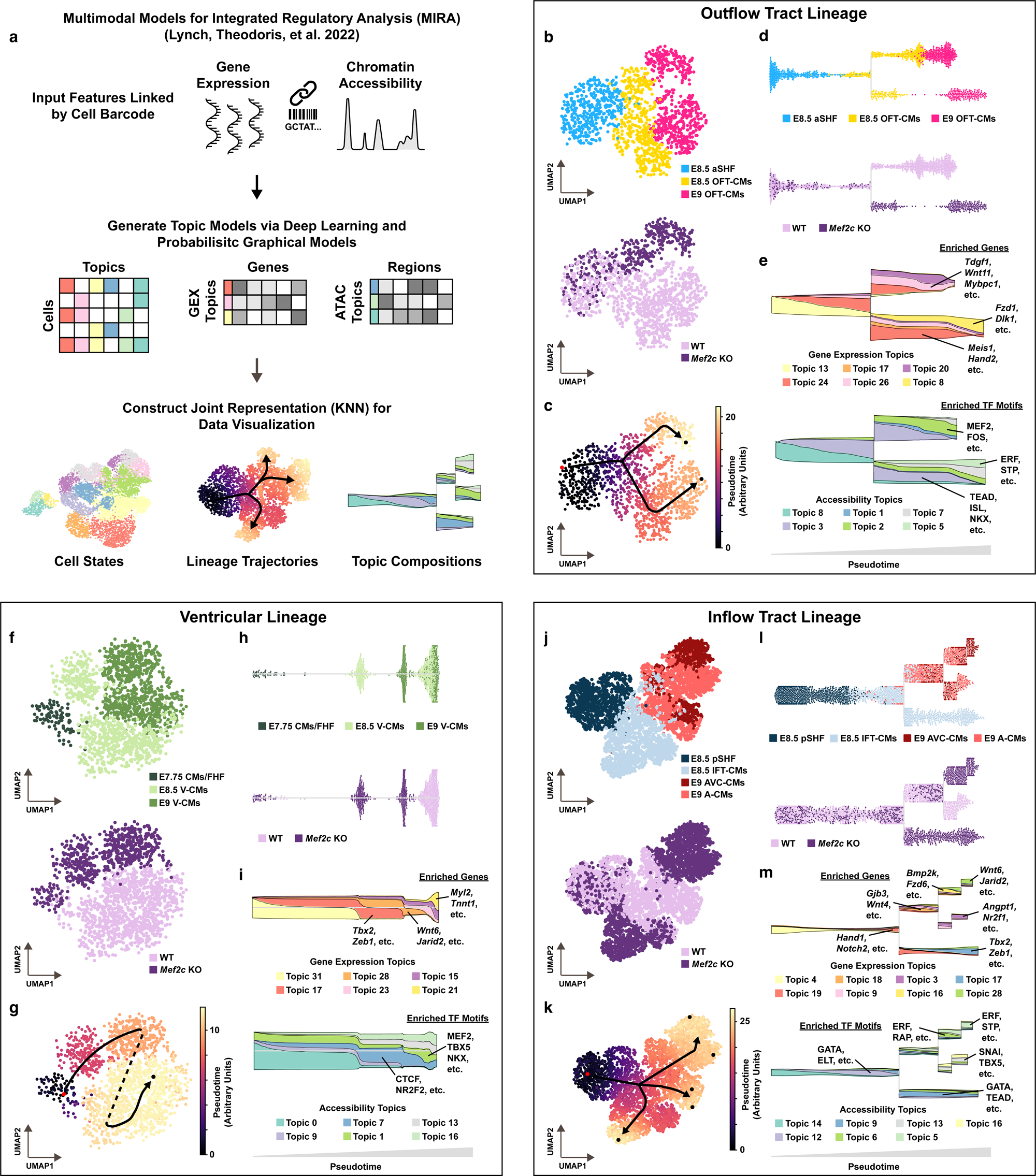
Each heart tube segment exhibits a distinct MEF2C-depedent developmental trajectory. a) Schematic of multimodal Models for Integrated Regulatory Analysis (MIRA)^24^ pipeline. b) UMAP of outflow tract lineage cells plotted by MIRA topic models labeled by cell type (top) and genotype (bottom). c) Pseudotime plot of outflow tract lineage cells. d) Lineage trajectory stream plots for outflow tract lineage cells labeled by cell type (top) and genotype (bottom). e) Stream plots displaying the flow of gene expression (top) and chromatin accessibility (bottom) topics in the outflow tract lineage cells. Examples of the top genes and TF binding motifs for dynamic topics are labeled. f-i) Same as (b-e), but for ventricular lineage cells. j-m) Same as (b-e) and (f-i), but for inflow tract lineage cells.

We trained topic models using cardiac progenitors and cardiomyocyte subtypes from both genotypes at each of our three timepoints (E7.75: CMs/FHF, SHF, JCF; E8.5: aSHF, pSHF, OFT-CMs, V-CMs, IFT-CMs; E9: OFT-CMs, V-CMs, A-CMs, AVC-CMs) and then used those models to construct joint representation UMAPs of the outflow tract, ventricular, and inflow tract lineages (Fig. 4b, 4f, 4j). We next calculated pseudotime values for cells within the UMAPs and used the results to construct cell state trajectories for each of the three lineages (Fig. 4c-d, 4g-h, 4k-l).

We observed clear differences in the fate specification of *Mef2c* KO cells compared to WT cells, as represented by branch points in the trajectories (Fig. 4d, 4h, 4l). Most interestingly, we noticed that the structure of these trajectories, along with the manner and timing by which *Mef2c* KO cells diverged from WT cells within the trajectories, was different for all three lineages. For instance, within the outflow tract lineage, *Mef2c* KO and WT cells proceeded similarly up until E8.5, and then diverged as development proceeded to E9 (Fig. 4d). Within the ventricular lineage, there was no branch point. Instead, *Mef2c* KO V-CMs from both E8.5 and E9 were located earlier in the trajectory than even E8.5 WT V-CMs (Fig. 4h). This indicates that loss of MEF2C in V-CMs resulted in a marked delay or termination of normal development rather than a divergence to an alternative cell state. The situation was most complex for the inflow tract lineage, in which there were two points of divergence between *Mef2c* KO and WT cells (Fig. 4l). First, a subset of *Mef2c* KO cells branched away from the WT trajectory as they differentiated from posterior SHF (pSHF) progenitors to IFT-CMs at E8.5. A second set of *Mef2c* KO cells proceeded along the WT trajectory to E9, before ultimately diverging from WT cells at the “terminal” AVC-CM and A-CM states. This complex branching behavior may reflect the notion that both FHF and pSHF progenitors contribute to the IFT^11^ and suggests that MEF2C likely controls distinct GRNs within these two IFT progenitor subpopulations. Together, these data reinforce our working hypothesis that MEF2C regulates development of the heart tube in a segment-specific manner via distinct regulatory networks.

We gained further insights regarding the segment-specific regulation of heart tube development by examining the top 200 genes and the top ranked TF binding motifs enriched in the gene expression and chromatin accessibility topics (Supplementary Table 5). As WT cells diverged from *Mef2c* KO cells in the outflow tract lineage, their gene expression topics became enriched for canonical outflow tract markers *Tdgf1* and *Wnt11*^21,25^, and their accessibility topics for MEF2 and FOS binding motifs (Fig. 4e). By contrast, the *Mef2c* KO cells were marked by gene expression topics enriched for Notch and Wnt signaling components such as *Dlk1* and *Fzd1*, and transcription factors *Meis1* and *Hand2,* and exhibited enrichment of an accessibility topic containing ISL, NKX, and TEAD binding motifs (Fig. 4e), suggesting these cells may retain a more progenitor-like state in the absence of MEF2C. The gene expression and accessibility topics enriched along the ventricular lineage trajectory supports our interpretation that the *Mef2c* KO cells experience a substantial delay or termination of their normal differentiation. Notably, topics that were enriched for CM genes (e.g. *Myl2*, *Tnnt1*) and cardiac TF binding motifs (e.g. TBX, MEF2, NKX) marked the terminal end of the differentiation trajectory consisting of only WT cells (Fig. 4i). Interestingly, the mid-point of the ventricular trajectory containing E8.5 and E9 *Mef2c* KO cells is marked by an accessibility topic enriched for the nuclear receptor NR2F2 binding motif, a transcription factor that is restricted to the IFT in WT embryos^26^ (Fig. 4i). In the complex branching inflow tract trajectory, terminal WT cells were enriched for topics containing expected atrial genes, such as *Angpt1* and *Nr2f1*, and binding motifs for TBX5 and SNAI, among others (Fig. 4m, Extended Data Fig. 4a-b). By contrast, the *Mef2c* KO cells streamed into one of two terminal branches enriched for genes such as *Wnt6, Jarid2, Tbx2,* and *Zeb1* and binding motifs for ERF, GATA, and TEAD factors (Fig. 4m, Extended Data Fig. 4a-b). Together, these data illustrate the lineage-specific changes in gene expression and chromatin accessibility that accompany heart tube development and the distinct impact of the loss of MEF2C on these lineage-specific programs.

### Candidate regulatory elements with MEF2C-dependent chromatin accessibility display enhancer activity in zebrafish

Our integrated chromatin accessibility data (Fig. 3) and developmental trajectory analyses with MIRA (Fig. 4) highlighted changes in chromatin accessibility during heart tube development that were MEF2C-dependent and segment-specific. We hypothesized that these regions contain regulatory elements that are necessary for driving segment-specific gene expression. To identify enhancers, we applied a series of filtering criteria to our integrated multiomic data and selected 12 candidates of the highest priority to screen based on the Log2FC value of peak accessibility and linkage to genes of interest (Fig. 5a, Supplementary Table 6). We enumerated these candidate regions with the prefix “MVEB,” after the initials of authors Muncie-Vasic and Brower. Overlaying our chromatin accessibility data at these regions with published ChIP-seq datasets^20,27^ confirmed that each of our candidates had clear and strong occupancy of both MEF2C and H3K27ac, a histone post-translational modification that marks active enhancers (Fig. 5b). Of note, one of our ventricular-specific candidates, MVEB8, was located within the *Nppa*/*Nppb* super enhancer cluster^28^. Furthermore, two of the candidates we identified from differential accessibility in IFT-CMs overlapped with regions found in the VISTA enhancer database^29^, with demonstrated enhancer activity in the hearts of E11.5 mouse embryos (Fig. 5c-d).

**Figure 5:**
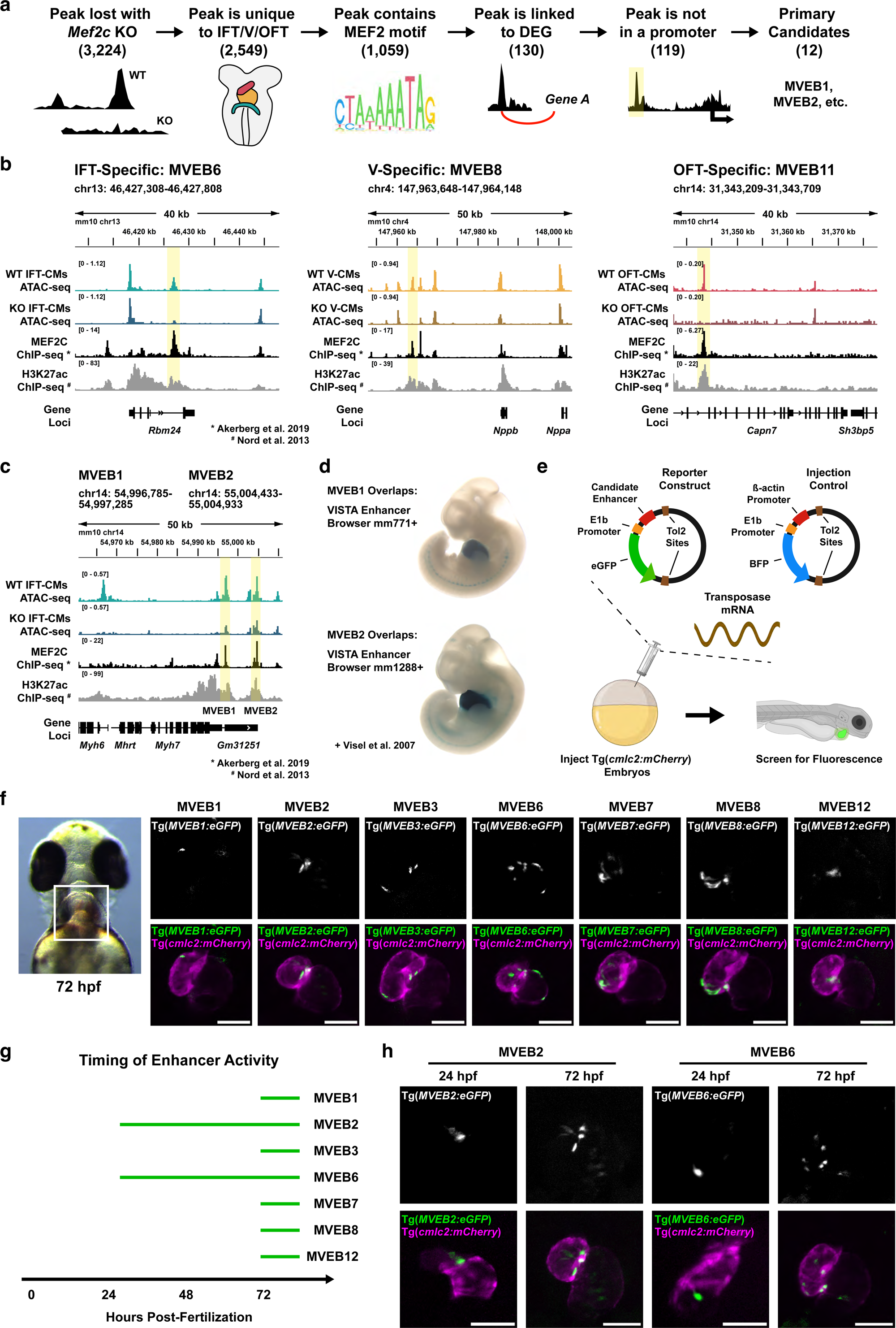
Candidate regulatory elements with MEF2C-dependent chromatin accessibility display enhancer activity in zebrafish. a) Schematic of selection process to identify candidate MEFC-dependent enhancers from integrated snRNA-seq and snATAC-seq data. b) Genome browser tracks displaying snATAC-seq accessibility profiles, MEF2C ChIP-seq occupancy profiles^20^, and H3K27ac ChIP-seq occupancy profiles^27^ at example loci containing IFT-specific (left), V-specific (middle), and OFT-specific (right) candidate enhancers (yellow highlights). c) Genome browser tracks displaying snATAC-seq accessibility profiles, MEF2C ChIP-seq occupancy profiles^20^, and H3K27ac ChIP-seq occupancy profiles^27^ at the *Myh6*/*Myh7* locus, which contains two candidate enhancers with IFT-specific altered accessibility (yellow highlights, MVEB1 and MVEB2) that overlap with regions found in the VISTA Enhancer Browser database^29^. d) Images of E11.5 mouse embryos from the Vista Enhancer Browser database^29^ demonstrating positive enhancer activity of regions that overlap with candidates MVEB1 and MVEB2. e) Schematic of the Tol2 transgenesis assay used to screen candidate enhancers in zebrafish. Elements of this panel were created in BioRender. B, B. (2024) https://BioRender.com/h83r503. f) Representative ventral view images of Tg(*cmlc2:mCherry*) zebrafish embryos at 72 hours post-fertilization (hpf) injected with candidate enhancers that demonstrated positive activity in the heart. Boxed area in the representative brightfield image (left) indicates the anatomical region of interest captured in the fluorescent images. g) Schematic representation of the observed onset of enhancer activity for candidate enhancers that demonstrated positive activity in the heart. h) Representative ventral view images of Tg(*cmlc2:mCherry*) zebrafish embryos at 24 and 72 hpf injected with *MVEB2:eGFP* or *MVEB6:eGFP* reporter constructs. Scale bars = 100 μm.

To determine whether our candidate regions are *bona fide* enhancers with activity in early heart tube development, we employed Tol2 transgenesis in zebrafish^30,31^, a relatively high-throughput *in vivo* screening system (Fig. 5e). We reasoned that because *Mef2c* has orthologs in zebrafish (*mef2ca* and *mef2cb*), and because there is a high degree of overlap between the key cardiac transcriptional regulators in both species^32^, there should be concordance between the activity of our candidate regions in mouse and zebrafish. Indeed, specific regulatory elements have been shown to exhibit activity in the heart of both species^33–35^.

Upon screening, seven of eleven candidates displayed enhancer activity in developing zebrafish hearts (Fig. 5f, Extended Data Fig. 5a-b), with one of our candidates being excluded due to a highly repetitive “GT” sequence. The two candidates that overlapped with regions found in the VISTA enhancer database (MVEB1, MVEB2) exhibited activity in the heart by 72 hours post-fertilization (hpf) (Fig. 5f). Additionally, two candidates from our IFT-CM data (MVEB3 and MVEB6), two from our V-CM data (MVEB7 and MVEB8), and one from our OFT-CM data (MVEB12) demonstrated robust and specific activity in the zebrafish heart at 72 hpf (Fig. 5f, Extended Data Fig. 5a-b). To our knowledge, four of these latter candidates (MVEB3, MVEB6, MVEB7, and MVEB12) have not been previously reported and represent novel, MEF2C-dependent enhancers with activity in the developing heart. Interestingly, we also observed a range in the timing of enhancer activity in the embryonic zebrafish heart. Candidates MVEB2 and MVEB6 were clearly detectable as early as 24 hpf, while the others were not active until 72 hpf (Fig. 5g-h). This variation in the onset of enhancer activity could result from differences in the timing of accessibility of these regulatory elements, different sensitivity to MEF2C levels, or cooperation with other TFs or transcriptional regulators.

### Loss of MEF2C induces an overactive posteriorized gene regulatory network driven by NR2F2 and GATA4

Our data indicate that loss of MEF2C causes a posteriorized gene signature (Fig. 2h-j) and chromatin accessibility landscape (Fig. 3d, 3f) in the developing heart tube. Moreover, as previously described^36^, the expanded but aberrantly formed inflow tract that forms in *Mef2c* KO embryos (Fig. 1c) indicates that loss of MEF2C leads to a more complex heart tube phenotype than a simple failure of cardiac differentiation and growth. To determine what transcriptional regulators may drive the inflow tract malformation, we performed TF binding motif enrichment analysis on the regions of chromatin that gain accessibility in the *Mef2c* KO IFT-CMs at E8.5. We found that Nuclear Receptor (NR) and GATA motifs were among the most enriched (Fig. 6a). From this observation, we hypothesized that MEF2C co-binds certain regulatory elements with NR or GATA factors in the WT condition, and that these NR and GATA factors become free to bind different regulatory elements with loss of MEF2C. Consistent with this hypothesis, we found that nearly half of the sites of accessible chromatin that mark WT IFT-CMs and contain an NR or GATA motif also contain a MEF2 motif, whereas the vast majority of sites that gain accessibility in *Mef2c* KO and contain an NR or GATA motif do not (Fig. 6b, Extended Data Fig. 6a).

**Figure 6:**
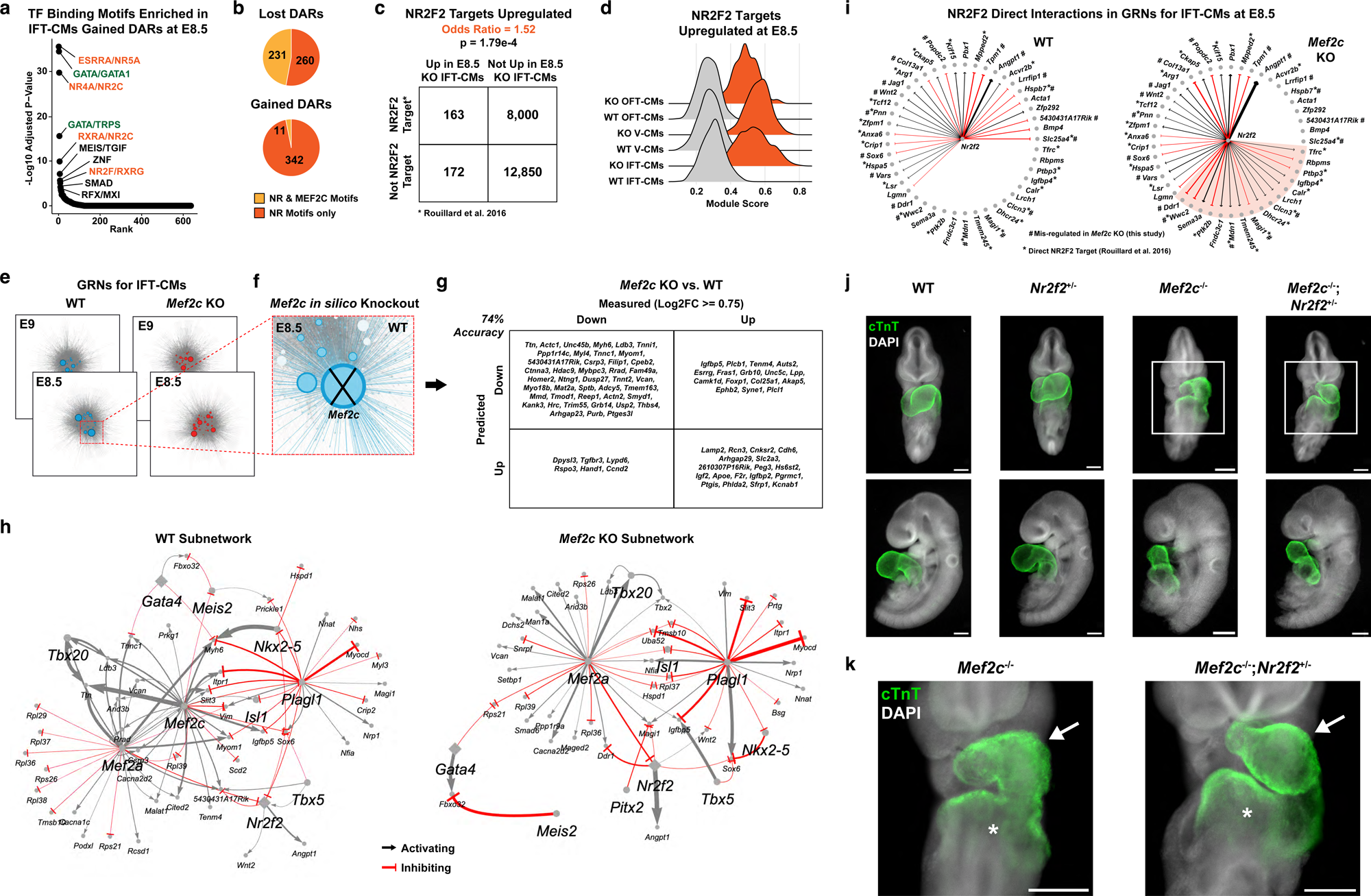
Loss of MEF2C induces an overactive posteriorized gene regulatory network that is partially rescued by reduced NR2F2 dosage. a) TF binding motif enrichment analysis for gained DARs in *Mef2c* KO relative to WT IFT-CMs at E8.5. b) Pie charts showing the proportion of lost or gained DARs (*Mef2c* KO-vs-WT) containing NR and MEF2C motifs or only NR motifs. c) Odds ratio analysis for NR2F2 target genes^40^ amongst DEGs up-regulated in *Mef2c* KO IFT-CMs. p-value calculated using Fisher’s exact test. d) Ridge plot displaying module scores for up-regulated NR2F2 targets in *Mef2c* KO and WT heart tube segments. e) Inferred GRNs constructed for *Mef2c* KO and WT IFT-CMs at E8.5 and E9. Boxed region of E8.5 WT GRN is shown at higher magnification in (f). f) Schematic of *in silico* simulated *Mef2c* KO in the E8.5 WT GRN. g) Results of GRN validation displaying the high accuracy (74%) of predicted relative to measured gene expression changes at E9. h) Visualizations of subnetworks consisting of 12 cardiac TFs and the top 100 DEGs within the WT and *Mef2c* KO E8.5 IFT-CM GRNs. i) Visualization of direct NR2F2 interactions in the WT and *Mef2c* KO E8.5 IFT-CM GRNs. Direct interactions that occur upon *Mef2c* KO are highlighted. #, mis-regulated DEG in *Mef2c* KO IFT-CMs at E8.5; *, Direct target of NR2F2^40^. j-k) Immunofluorescent staining of cardiac Troponin T (cTnT, green) in representative E9.5 embryos (18-24 somites) collected from *Mef2c*^+/-^;*Nr2f2*^+/-^ to *Mef2c*^+/-^ crosses. Boxed regions in (j) are shown at higher magnification in (k). Arrows point to the ventricle, which is expanded in *Mef2c*^-/-^;*Nr2f2*^+/-^ embryos compared to *Mef2c*^-/-^ embryos. Asterisks mark the atria, which are better developed and have undergone more looping in *Mef2c*^-/-^ ;*Nr2f2*^+/-^ embryos compared to *Mef2c*^-/-^ embryos. n=5 *Mef2c*^-/-^ and n=7 *Mef2c*^-/-^;*Nr2f2*^+/-^ embryos from 7 independent litters. Scale bars = 200 μm

These data are particularly interesting given the importance of GATA factors, and in particular GATA4, in cardiac development^37^ and the knowledge that nuclear receptor subfamily 2 group F member 2 (NR2F2, also known as COUP-TFII) is a necessary driver of atrial development^26,38^. Using publicly available ChIP-seq datasets^38,39^, we found NR2F2 and GATA4 occupancy at a number of the regions of chromatin that gained accessibility in the *Mef2c* KO IFT-CMs at E8.5 (Extended Data Fig. 6b-c), suggesting that these DARs could indeed be sites of potential NR2F2 and GATA4 regulation. To determine whether altered NR2F2 and GATA4 activity may drive the posteriorization of the heart tube, we next asked if NR2F2 and GATA4 target genes were upregulated in the inflow tract of *Mef2c* KO embryos. Using publicly available datasets to identify the target genes of each TF^40,41^, we calculated the odds ratio (OR) of finding NR2F2 or GATA4 target genes compared to non-target genes amongst our up-regulated DEGs at E8.5. We found strong evidence that NR2F2 (OR = 1.52, p = 1.79e-4) and GATA4 (OR = 4.23, p < 2.2e-16) targets were enriched amongst the upregulated genes (Fig. 6c, Extended Data Fig. 6d). Next, we took the list of NR2F2 and GATA4 target genes that were upregulated (Supplementary Table 7) and calculated a module score^42^ for the expression of these genes in our dataset. In agreement with our working model that these NR2F2 and GATA4 targets are driving the posteriorized heart tube phenotype in *Mef2c* KO, we found the NR2F2 and GATA4 module scores were increased in all segments of the heart tube at E8.5 (Fig. 6d, Extended Data Fig. 6e).

To understand how altered NR2F2 and GATA4 activity upon loss of MEF2C affects the entire transcriptional network, we used CellOracle^43^ to construct GRNs for the WT and *Mef2c* KO IFT-CMs at E8.5 and E9 (Fig. 6e). In brief, CellOracle uses integrated gene expression and chromatin accessibility data to infer interactions between TFs and target genes based on the accessibility of a given TF’s binding motif near a potential target gene and the correlation of expression between the TF and the potential target. To validate that these inferred CellOracle networks effectively captured the GRNs that govern inflow tract development, we simulated the knockout of *Mef2c in silico* in the E8.5 WT IFT-CMs network and compared the predicted changes in gene expression to the measured changes captured in our E9 dataset (Fig. 6f-g). We found that the networks correctly predicted the gene expression changes with 74% accuracy, even with a relatively permissive DEGs cutoff of Log2FC >= 0.75 (Fig. 6g). As expected, the prediction accuracy increased with more stringent Log2FC cutoffs, and vice versa (Supplementary Table 8).

We next visualized the subnetworks of 12 core cardiac TFs and the top 100 DEGs in the WT and *Mef2c* KO inflow tracts (Fig. 6h). These subnetworks revealed that loss of MEF2C causes a network-wide reorganization of gene regulation. First, we observed that MEF2C may be critical for coordinating cooperative regulation by multiple cardiac TFs. For instance, in the WT network, MEF2C, MEF2A, and TBX20 all have an inferred activating effect on *Ttn* expression. However, in the *Mef2c* KO network, both MEF2A and TBX20 lose their activating interaction with *Ttn.* Similarly, in the WT network, MEF2C, MEF2A, and NKX2-5 are all inferred activators of *Myh6*, but in the *Mef2c* KO network, MEF2A and NKX2-5 are no longer predicted to activate *Myh6* expression. These data suggest that MEF2C is required for cooperative activation with other cardiac TFs in the developing heart tube, similar to what was shown to occur during cardiomyocyte reprogramming *in vitro*^44^. Additionally, we found that loss of MEF2C may alter the scope and strength of interactions between other cardiac TFs and their gene targets. For example, in the WT network, GATA4 and MEIS2 exhibit competitive regulation of *Fbxo32*, which encodes a muscle-specific ubiquitin-E3 ligase that is critical for sarcomeric function and associated with dilated cardiomyopathy^45,46^. This competitive relationship is maintained in the *Mef2c* KO network, but the interactions are strengthened in both directions. Finally, we observed that the activity of NR2F2 is substantially altered in the *Mef2c* KO network, with a larger number of gene targets and apparent strengthening of both activating and inhibitory interactions. Most notably, the activating interaction between NR2F2 and *Angpt*1 expression is much stronger in the *Mef2c* KO network than in the WT network. Interestingly, the activation of *Nr2f2* itself by TBX5 is lost upon *Mef2c* KO.

Next, we further narrowed in on the direct interactions between NR2F2 and GATA4 and their target genes in the WT and *Mef2c* KO inflow tract networks (Fig. 6i, Extended Data Fig. 6f). From these analyses we observe similar behavior for both TFs – approximately half of the WT interactions are maintained in the *Mef2c* KO, while nearly twice as many new direct interactions are established. This is consistent with our working model that in response to loss of MEF2C, aberrant NR2F2 and GATA4 activity regulate additional genes that drive the inflow tract phenotype in the posterior of the heart tube.

### Reduction of NR2F2 dosage partially rescues the heart tube phenotype in *Mef2c* **KO embryos**

Given our observation that loss of MEF2C causes aberrant NR2F2 activity in the IFT-CM GRNs, we theorized that reducing the dosage of NR2F2 may rescue the inflow tract phenotype in *Mef2c* KO embryos. To test this possibility, we generated *Mef2c*^+/-^;*Nr2f2*^+/-^ male mice and crossed them to *Mef2c*^+/-^ females. Although *Nr2f2*^+/-^ embryos have no apparent heart tube phenotype at E9.5 (Fig. 6j, Extended Data Fig. 7a), pups of this genotype were recovered at lower numbers than expected Mendelian ratios^26^, suggesting that *Nr2f2* is a dosage-sensitive gene. We observed a notable partial rescue in ventricle and inflow tract development at E9.5 (18-24 somites) in the *Mef2c*^-/-^;*Nr2f2*^+/-^ embryos compared to *Mef2c*^-/-^ (Fig. 6j-k), although the extent of the rescue varied (Extended Data Fig. 7b-c). The *Mef2c* KO single-ventricle phenotype persisted in *Mef2c^-/-^*;*Nr2f2*^+/-^ embryos, but these ventricles were larger and more expanded than in *Mef2c^-/-^ embryos.* Moreover, the posterior structures of the heart tube in *Mef2c^-/-^*;*Nr2f2*^+/-^ embryos more closely resembled WT inflow tracts with expanded and looping prospective atria, compared to the completely disrupted inflow tract in *Mef2c*^-/-^ embryos (Fig. 6j-k). This partial rescue of the ventricle and inflow tract in *Mef2c^-/-^*;*Nr2f2*^+/-^ embryos is evidence that the *Mef2c* KO phenotype in the posterior heart tube is driven, at least in part, by an altered gene regulatory network with increased and aberrant NR2F2 activity.

## DISCUSSION

In this study we demonstrated that MEF2C controls complex, segment-specific GRNs that regulate gene expression and chromatin accessibility in the developing linear heart tube. We found that loss of MEF2C causes a posteriorization of the heart tube and revealed novel segment-specific regulatory elements with enhancer activity. These results address the long-standing question of how a given TF can play distinct regulatory roles in different cell types and at particular developmental stages. For instance, decades of work have established that FHF progenitors give rise to the left ventricle and atria, while the SHF is subdivided into the aSHF, which produces the outflow tract and right ventricle, and the pSHF, which contributes to the atria^7–11^. Although these progenitor cell types have different molecular signatures and give rise to individual structures with specific morphology and function, they largely rely on a shared set of cardiac TFs to regulate their development. Our data illustrate that these core cardiac TFs operate in unique GRNs for each of the inflow tract, ventricular, and outflow tract lineages in the developing heart tube. We found that distinct chromatin landscapes and regulatory elements are important determinants for setting up these GRNs, but other embryological phenomena are likely to be involved, such as morphogen signaling from differing neighbor cell types, heterochronic timing of differentiation^47^, and mechanical phenomena introduced by tissue morphogenesis and the initiation of cardiomyocyte beating^48–50^.

Using the GRNs inferred from our multiomic data, we identified a genetic interaction between *Mef2c* and *Nr2f2* in the inflow tract of the developing heart tube. Previous work from our lab showed that TF occupancy at specific loci is highly dependent upon both TF-TF and TF-DNA affinities, and that TF-TF interactions not only serve to regulate gene expression, but prevent the redistribution of one or both TFs to exogenous loci^51^. The results we present here lend further support to those concepts. By removing MEF2C, the set of possible TF-TF interactions is limited and causes altered activity of partner TFs, including NR2F2 and GATA4. Our comparison of the WT and *Mef2c* KO GRNs revealed that the transcriptional networks regulating early heart development are highly sensitive to TF perturbations, with loss of MEF2C causing a reorganization of heart tube GRNs that is not limited to direct MEF2C connections, and reduction of NR2F2 dosage capable of partially rescuing the heart tube malformations. Thus, these GRNs may be implemented as a powerful tool for predicting the network-wide consequences of genetic mutations that underlie CHDs, potentially leading to unexpected molecular targets for intervention and correction.

Our approach provides a generalizable framework for integrating snRNA-seq and snATAC-seq data to build GRNs that provide specific and comprehensive models of gene regulation. By constructing these GRNs using both WT and mutant data, we were able to immediately test predictions made from the WT networks and validated the inferred regulatory interactions. These tools also allow specific networks to be built for particular cell or tissue types, provided there are enough cells in the group of interest to generate robust inferences. Moreover, the experimental mutant data revealed how the overall networks were altered – both in ways that would and would not be predicted from the WT networks. These unexpected differences help to illuminate indirect or transactivating gene regulation. Discoveries from these networks lead to many new hypotheses about regulatory elements, TF interactions, DNA binding, and chromatin organization. As these hypotheses are experimentally tested, the GRNs can be refined, leading to higher-fidelity models with greater predictive power. Such higher-fidelity networks will yield important new insights into the transcriptional regulation of developmental processes and will have the ability to reveal precisely how these processes are altered in the context of disease-causing mutations.

## MATERIALS AND METHODS

### Mouse Models

All mouse studies were performed in strict compliance with the UCSF Institutional Animal Care and Use Committee. Mice were housed in a standard 12-hour light/dark animal husbandry barrier facility at the Gladstone Institutes. The *Mef2c*^+/-^ allele used was the same as the original generated by Lin et al.^13^. The *Smarcd3*-F6-eGFP reporter allele was generated in our lab and described previously^14^. The *COUP-TFII lacZ* knock-in allele (abbreviated here as *Nr2f2*^+/-^) was obtained from the Mutant Mouse Resource & Research Centers and was originally generated by Takamoto et al.^52^. The *Mef2c*^+/-^ and *Nr2f2*^+/-^ lines were maintained in the C57BL6/J background (Jackson Laboratory #664). The *Smarcd3*-F6-eGFP allele was crossed into our *Mef2c*^+/-^ line from a mixed background of C57BL6/J (Jackson Laboratory #664) and CD-1 (Charles River #022).

### Timed Embryo Dissections and Genotyping

To achieve timed matings, male and female mice were housed together in the evening and pregnancy was assessed by vaginal plug the following morning. Gestational stage was determined relative to noon on the day of plug detection, defined as day E0.5. Embryos were dissected and, at later stages when yolk was present, also de-yolked, in ice-cold PBS (Life Technologies, 14190250) with 1% fetal bovine serum (FBS; Thermo Fisher Scientific, 10439016) on ice. The posterior trunk was removed by microdissection for all embryos at E8.5 or later that were collected for imaging (IF or FISH via HCR) to ensure unobstructed views of the heart tube. DNA for genotyping was extracted using QuickExtract DNA Extraction Solution (Lucigen, QE09050) from harvested yolk sac tissue, if available, or else from a micro-dissected piece of the extra-embryonic anterior proximal region. For embryos collected for 10x Multiome library construction, rapid genotyping was performed while embryos remained on ice using Phire Green Hot Start II PCR Master Mix (Thermo Fisher Scientific, F126S), according to manufacturer’s protocols. For all other experiments, genotyping was performed after embryo fixation using GoTaq Green Master Mix (VWR, PAM7122), according to manufacturer’s protocols. PCR products were run on a 1% agarose gel (Thermo Fisher Scientific, BP164-500) with Biotium GelRed Nucleic Acid Gel Stain (Thermo Fisher Scientific, NC9594719) used for DNA detection. Primers used for genotyping were ordered from Integrated DNA Technologies (IDT). For genotyping the *Mef2c* allele, a three-primer mix was used (P1: ACTGGCTGAGAGTTGTACCCAC, P2: GCGTGATTTGCTAGATAGTGGTAGAC, P3: ATGTGGAATGTGTGCGAGGC), resulting in a single 314 bp band for WT, a single 242 bp band for *Mef2c*^-/-^, and both bands for *Mef2c*^+/-^. For genotyping the *Nr2f2* allele, a three-primer mix was used (P4: CATCCGGGATATGTTACTGTCCGG, P5: TGGGGAAGCTAAGTGTTGATGTGATTCC, P6: GCCGTGGGTTTCAATATTGGCTTC), resulting in a single 786 bp band for WT, a single ∼1600 bp band for *Nr2f2*^-/-^, and both bands for *Nr2f2*^+/-^.

### Whole-Mount Embryo Immunostaining and Imaging

Embryos collected for immunostaining were fixed for 1 h at room temperature in 4% paraformaldehyde (PFA) freshly diluted from 16% weight/volume PFA aqueous solution (Thermo Fisher Scientific, 043368-9M) in PBS (Life Technologies, 14190250). Embryos were stored at 4°C for up to 1 month prior to immunostaining. To permeabilize and block non-specific antigens, embryos were incubated at 37°C for 2 h with gentle rotation in a blocking buffer consisting of PBS (Life Technologies, 14190250), 5% normal donkey serum (Sigma Aldrich, S30-M), and 0.8% Triton X-100 (Sigma Aldrich, X-100-5ML). This was then removed and replaced with primary antibodies diluted in the same blocking buffer and incubated overnight at 37°C with gentle rotation. The next day, primary antibody solution was removed and embryos were washed 3x 30 min in the same blocking buffer at 37°C with gentle rotation. Secondary antibodies and DAPI (Abcam, ab228549, used at 1 μg/mL) were added in a blocking buffer consisting of PBS (Life Technologies, 14190250), 5% normal donkey serum (Sigma Aldrich, S30-M), and 0.4% Triton X-100 (Sigma Aldrich, X-100-5ML), and incubated for 3-4 h at 37°C with gentle rotation. Secondary antibody solution was removed and embryos were washed 3x 30 min in PBS (Life Technologies, 14190250) with 0.1% Tween-20 (Thermo Fisher Scientific, J20605.AP) at 37°C with gentle rotation. Embryos were imaged in PBS using an upright epifluorescence microscope (Leica MZFLIII, Leica DFC 3000G, Lumen Dynamics XCite 120LED) and acquisition software LASX (Leica). Primary antibodies used were sheep polyclonal MEF2C (R&D Systems, AF6786, used at 1:250) and rabbit polyclonal cardiac Troponin T (Thermo Fisher Scientific, 15513-1-AP, used at 1:250). Secondary antibodies used were Donkey anti-Sheep IgG (H+L) Cross-Adsorbed Secondary Antibody, Alexa Fluor™ 647 (Thermo Fisher Scientific, A-21448, used at 1:1000) and Donkey anti-Rabbit IgG (H+L) Highly Cross-Adsorbed Secondary Antibody, Alexa Fluor™ 555 (Thermo Fisher Scientific, A-31572, used at 1:1000).

### Whole-Mount Embryo Fluorescence *In Situ* Hybridization via Hybridization Chain Reaction and Imaging

The protocol for *in situ* hybridization via hybridization chain reaction (HCR™) was adapted from the optimized manufacturer’s protocol for whole-mount mouse embryos (Molecular Instruments). Embryos were fixed overnight at 4°C in 4% PFA freshly diluted from 16% weight/volume PFA aqueous solution (Thermo Fisher Scientific, 043368-9M) in PBS (Life Technologies, 14190250). Embryos were then washed 3x 5 min in PBS (Life Technologies, 14190250) with 0.1% Tween-20 (Thermo Fisher Scientific, J20605.AP), abbreviated PBST, and then processed through 10 min serial incubations in a dehydration series of 25%, 50%, 75%, and 100% methanol solutions in PBST on ice. Embryos were stored in 100% methanol at −20°C for up to 6 months prior to *in situ* hybridization. Embryos were then rehydrated through 10 min serial incubations in a series of 75%, 50%, 25%, and 0% methanol solutions in PBST on ice. Embryos were then brought to room temperature and permeabilized by 10 min incubation with 10 μg/mL Proteinase K (Sigma Aldrich, 70663-5) in PBST, washed 2x 5 min in PBST at room temperature, post-fixed in 4% PFA freshly diluted from 16% weight/volume PFA aqueous solution (Thermo Fisher Scientific, 043368-9M) in PBS (Life Technologies, 14190250), and then washed 3x 5 min in PBST at room temperature. Embryos were pre-hybridized in Probe Hybridization Buffer (Molecular Instruments) for 5 min at room temperature followed by an additional 30 min at 37°C with fresh buffer. Single or multiplexed hybridization probes were then diluted in pre-warmed Probe Hybridization Buffer (Molecular Instruments) and embryos were hybridized for 48 h at 37°C. Embryos were then washed 4x 15 min with pre-warmed Probe Wash Buffer (Molecular Instruments) at 37°C, followed by 2x 5 min with 5xSSCT (diluted in nuclease-free water from 20xSSCT, Thermo Fisher Scientific, 15557044) at room temperature. During washes, amplification hairpins were snap-cooled by heating to 95°C for 90 s and then allowed to cool to room temperature in the dark. Embryos were pre-amplified in Amplification Buffer (Molecular Instruments) for 5 min at room temperature. Snap-cooled hairpins and DAPI (Abcam, ab228549, used at 1 μg/mL) were diluted in Amplification Buffer (Molecular Instruments) and embryos were incubated in this solution overnight in the dark at room temperature. The following day, embryos were washed through a series of 5xSSCT washes (diluted in nuclease-free water from 20xSSCT, Thermo Fisher Scientific, 15557044) in the dark at room temperature (2x 5 min, 2x 30 min, 1x 5 min). Embryos were imaged in PBS using an upright epifluorescence microscope (Leica MZFLIII, Leica DFC 3000G, Lumen Dynamics XCite 120LED) and acquisition software LASX (Leica). To maximize the utility of collected embryos, probes were then stripped by overnight incubation in 80% formamide in water (Thermo Fisher Scientific, BP228100) at room temperature, followed by 3x 5 min washes in PBST at room temperature. Embryos could then be probed for additional targets, beginning the protocol again at the pre-hybridization step. Probes used in this study were ordered from the Molecular Instruments catalog, when possible, or custom-designed and manufactured by Molecular Instruments. Hybridization probes used were: *Tnnt2*-B3 (used at 4 nM), *Ttn*-B2 (used at 4 nM), *Nppa*-B2 (used at 20 nM), *Nkx2.5*-B1 (used at 20 nM), *Tbx5*-B1 (used at 20 nM), *Gata4*-B1 (used at 20 nM), *Wnt2*-B2 (used at 20 nM), *Tdgf1*-B2 (used at 20 nM). Amplification probes used were all used at 60 nM: B1-Alexa Fluor™ 647, B2-Alexa Fluor™ 546, B2-Alexa Fluor™ 647, B3-Alexa Fluor™ 488.

### Embryo Preparation for 10x Multiome snRNA-seq and snATAC-seq

For E7.75 embryos, whole embryos were dissected and harvested for single-nucleus library generation. For E8.5 and E9 embryos, the head folds and posterior trunk were removed by microdissection prior to harvesting for library generation to enrich the relative capture of cardiac cell types. Two embryos were collected per genotype per embryonic stage. Following dissection and rapid PCR genotyping, embryo samples selected for the multiome experiment were incubated in 200 μl TrypLE Select (Thermo Fisher Scientific, 12563-011) for 5 min at 37°C, triturated gently by pipetting up and down, and then incubated an additional 3 min at 37°C. The dissociated cell suspension was quenched with 600 μl of PBS (Life Technologies, 14190250) with 1% FBS (Thermo Fisher Scientific, 10439016), singularized by passage through the 35 μm mesh of a 5 mL Falcon™ Round-Bottom Polystyrene Test Tube with Cell Strainer Snap Cap (Thermo Fisher Scientific, 08-771-23), pelleted by centrifugation at 300 g for 5 min at 4°C, and resuspended in 50 μl of PBS (Life Technologies, 14190250) with 0.04% bovine serum albumin (BSA, Sigma Aldrich, A1595). At this stage, the manufacturer’s protocol for Nuclei Isolation for Single Cell Multiome ATAC + Gene Expression Sequencing, Appendix, Low Cell Input Nuclei Isolation (10x Genomics, CG000365 Rev B) was followed exactly to prepare nuclei for library generation.

### Generation of snRNA-seq and snATAC-seq Libraries and Sequencing

Libraries for snRNA-seq and snATAC-seq were prepared according to the manufacturer’s protocol for Chromium Next GEM Single Cell Multiome ATAC + Gene Expression (10x Genomics, CG000338 Rev B), using the 10x Genomics Chromium controller, Chromium Next GEM Single Cell Multiome ATAC + Gene Expression Reagent Bundle (10x Genomics, PN-1000283), Chromium Next GEM Chip J Single Cell Kit (10x Genomics, PN-1000234), Single Index Kit N Set A (10x Genomics, PN-1000212), and Dual Index Kit TT Set A (10x Genomics, PN-1000215). Nuclei isolated from multiome samples were processed directly into the snATAC-seq transposition reaction and then captured in 10x Genomics GEMs via the Chromium Controller. GEMs were stored at -80°C for up to 4 weeks, allowing for collection of replicate embryos of each genotype at each development stage. All subsequent library preparation steps were performed together for all embryos of a given developmental stage to reduce the likelihood of batch artifacts. A targeted maximum recovery of 10,000 cells per sample were loaded onto the 10x Genomics Chromium controller instrument and each sample was indexed with a unique sample identifier (Single Index Kit N Set A for snATAC-seq libraries, Dual Index Kit TT Set A for snRNA-seq libraries). Final libraries were quality-controlled using an Agilent Bioanlyzer instrument with Agilent High Sensitivity DNA Kit (Agilent, 5067-4626). The DNA concentration of each library was measured using KAPA Library Quantification qPCR Kit (Thermo Fisher Scientific, 50-196-5234), and then libraries were pooled and sequenced on NovaSeq6000 S4 lanes (Illumina). All snRNA-seq libraries were sequenced to a depth of at least 20,000 mean raw aligned reads per cell and snATAC-seq libraries were sequenced to a depth of at least 35,000 mean raw aligned read pairs per cell (most >60,000).

### Processing of Raw Sequencing Data

Raw sequencing reads were processed using the 10x Genomics Cell Ranger Arc v2.0.0 pipeline. Reads were demultiplexed using cellranger-arc mkfastq and aligned with cellranger-arc count to the mm10 reference genome containing an additional sequence for eGFP to map reads of the *Smarcd3*-F6-eGFP reporter transcript.

### Seurat Analysis of snRNA-seq Data

Gene expression data were analyzed using Seurat v4.3.0^15^ in R. Datasets for each developmental stage (E7.75, E8.5, E9) were analyzed as separate Seurat objects. Counts matrices from each sample that were output by the Cell Ranger Arc pipeline were used as inputs to the Read10X() and CreateSeuratObject() functions. Data were filtered for quality control based on number of detected genes, unique molecular identifier (UMI) counts, percent ribosomal transcripts, and percent mitochondrial transcripts. SCTransform v2^16^ was used to normalize and scale data, with regressions performed with respect to percentage of ribosomal transcripts and percentage of mitochondrial transcripts. Principal component analysis (PCA) was performed, followed by batch correction with Harmony^53^, and cells were clustered and visualized as UMAP embeddings based on the top 30 principal components using FindNeighbors(), FindClusters(), RunUMAP(), and DimPlot(). Cell types were annotated using the FindAllMarkers() function with Wilcoxon rank-sum test (min.pct = 0.25, logfc threshold = 0.25, p_val_adj < 0.05) to identify cluster-specific marker genes (Supplementary Table 1, Extended Data Fig. 2d-f). Cardiac progenitors, cardiomyocytes, and related mesoderm cell types were subset into new Seurat objects. PCA analysis, batch correction, visualization, and clustering were then performed on these new objects as described above. Cell types in the subset objects were annotated using the FindAllMarkers() function with Wilcoxon rank-sum test (min.pct = 0.25, logfc threshold = 0.5, p_val_adj < 0.05) to identify cluster-specific marker genes that enabled more specific cell type labeling than in the full datasets (Supplementary Table 2, Extended Data Fig. 2g-i). Differential gene expression was performed between *Mef2c* KO and WT cell types of interest using the FindMarkers() function with Wilcoxon rank-sum test (min.pct = 0.1, logfc threshold = 0.25, p_val_adj < 0.05). The DotPlot() and RidgePlot() functions was used to visualize gene expression levels of differentially expressed genes across cell types of interest. The AddModuleScore() function(42) was used to add module scores for up-regulated (*Mef2c* KO-vs-WT) NR2F2 and GATA4 target genes and the RidgePlot() function was used to visualize these module scores in cell types of interest.

### ArchR Analysis of Integrated snRNA-seq and snATAC-seq Data

#### Initial Dataset Processing

Chromatin accessibility and integrated multiome data was analyzed using the ArchR software package v1.0.2^22^ in R. The fragments files output by the Cell Ranger Arc pipeline were used as inupts to the createArrowFiles() function to create Arrow Files for each sample. These Arrow Files were used to create individual ArchR projects for developmental stage (E7.75, E8.5, E9). The gene expression data was added using the import10xFeatureMatrix() and addGeneExpressionMatrix() functions, using the matching cell barcodes to combine the snRNA-seq and snATAC-seq data for each cell. Data were filtered for quality control based on the number of unique fragments, TSS Enrichment score, expression UMI counts, and number of detected genes. Doublets were filtered using the addDoubletScores() and filterDoublets() functions. Dimensionality reduction was performed using ArchR’s iterative latent semantic index (LSI) implementation individually for the snATAC-seq data (varFeatures = 25,000, dimsToUse = 1:30) and snRNA-seq data (varFeatures = 5,000, dimsToUse = 1:30). Then, the reduced dimensions were combined using addCombinedDims() and batch correction was performed on the combined dimensions using Harmony^53^. Cells were then embedded in UMAP space with addUMAP(), clustered using addClusters(), and visualized using plotEmbedding(). Cell types were annotated by transferring metadata labels from the previously analyzed Seurat objects. The getMarkerFeatures() function with Wilcoxon rank-sum test (FDR <= 0.01, Log2FC >= 0.5) was used to identify cluster-specific marker genes to confirm cell type labels. Cardiac progenitors, cardiomyocytes, and related mesoderm cell types were subset into new ArchR projects. Dimensionality reduction, batch correction, clustering, and visualization were performed as described above. Cell types in the subset projects were annotated by transferring metadata labels from the previously analyzed subset Seurat objects. The getMarkerFeatures() function with Wilcoxon rank-sum test (FDR <= 0.01, Log2FC >= 0.5) was used to identify cluster-specific marker genes to confirm subset cell type labels. Due to the small number of *Mef2c* KO OFT-CMs in the E8.5 dataset, an ArchR project with combined E8.5 and E9 samples was created and processed as described above for downstream analyses/comparisons of OFT-CMs.

### Peak Calling, Differential Peak Testing, and Motif Enrichment

Cells were pseudobulked into replicates based on their cell type and genotype labels (e.g. WT V-CMs), with two pseudobulked replicates created corresponding to the two embryos collected for each genotype at each developmental stage. Replicates were created using the addGroupCoverages() function (minCells = 20, maxCells = 500, sampleRatio = 0.8). Peaks were then called using addReproduciblePeakSet() with MACS2(23) (reproducibility = 2, peaksPerCell = 500, minCells = 20). Differential peak accessibility testing was performed using getMarkerFeatures() with Wilcoxon rank-sum test (FDR <= 0.15, Log2FC >= 0.5 or Log2FC <= -0.5) for each cell type of interest with *Mef2c* KO cells used as the group of interest and WT cells used as the background. Transcription factor binding motifs were annotated using addMotifAnnotations() with the Vierstra motif archetypes^54^ and motif enrichments for regions of gained or lost accessibility were calculated using peakAnnoEnrichment() with cutoffs of FDR <= 0.15 and Log2FC <= -0.5 or Log2FC >= 0.5.

### Peak2Gene Linkages and Browser Track Visualization

Peak2Gene Linkages were identified using the addPeak2GeneLinks() and getPeak2GeneLinks() functions. Code written in this study was used to filter Peak2Gene links based on presence of a MEF2C motif and linkage to specific genes (i.e. DEGs). Peak2Gene links were visualized using the plotBrowserTrack() function. The getGroupBW() function was used to export the accessibility coverage data in bigwig format for visualization along with published ChIP-seq data using the IGV software package^55^.

### MIRA Analysis of Integrated snRNA-seq and snATAC-seq Data

#### Preparing Merged Datasets

We used the Multimodal models for Integrated Regulatory Analysis (MIRA)^24^ python package to examine dynamic changes in gene expression and chromatin accessibility in WT and *Mef2c* KO embryos across the time course of developmental stages that we collected (E7.75, E8.5, E9). We first created a merged Seurat object and ArchR project containing only our cardiac cell types of interest. We then used sceasy^56^ to convert our Seurat object and the peak matrix from the ArchR project to AnnData formats, which were used as inputs for the MIRA analyses.

### Topic Modeling

The first step of the MIRA pipeline is to train and tune topic models for both the gene expression and chromatin accessibility data. Model training and tuning was performed with GPU support on nodes of the Wynton HPC Co-Op cluster at UCSF. For our gene expression topic model, we used the 4,088 most variable genes (min_disp = 0.1) as the “exogenous” gene set and top 1,959 variable genes (dispersion threshold > 0.7) as the “endogenous” gene set for model training. We then set the minimum and maximum learning rate bounds, tuned the model (iters = 64, max_topics = 55), and MIRA’s tuner selected the best model that was used for all subsequent analyses, which contained 33 gene expression topics. To aid in the interpretation of results, we exported the top 200 genes for each of the 33 gene expression topics. We then repeated these steps to train and tune the accessibility topic model, using all features (i.e. there are no “exogenous” or “endogenous” peak sets), which resulted in a model containing 17 accessibility topics. To aid in the interpretation of these topics, we performed TF binding motif enrichment analysis on each topic and exported the TF motif enrichment scores (top_quantile = 0.2) for each of the 17 accessibility topics.

### Pseudotime Analysis and Lineage Trajectories

For each of the heart tube segment lineages that we analyzed (OFT, V, IFT), we created separate subsetted AnnData objects containing only the cell types that contribute to each lineage. We generated UMAP embeddings for each lineage based on joint representations of gene expression and accessibility topics. We then performed pseudotime analysis by calculating the diffusion map, selecting the appropriate number of diffusion map components, and generated a K nearest neighbor graph based on these diffusion map components. To create pseudotime trajectories, we randomly selected a WT cell in the earliest progenitor cell type of each lineage to serve as the initial cell, and then manually selected terminal cells from the appropriate cell types based on the structure of each trajectory (for OFT lineage: E9 WT OFT-CM and E9 KO OFT-CM; for V lineage: E9 WT V-CM; for IFT lineage: E8.5 KO IFT-CM, E9 WT A-CM, E9 WT AVC-CM, E9 KO A-CM, E9 KO AVC-CM). Given these initial and terminal cells, along with the KNN pseudotime graph, the probability of reaching each terminal state was calculated for every cell and these probabilities we parsed to create the bifurcating tree lineage trajectory plots (stream plots). Gene expression and accessibility topics of interest were then plotted along these stream plots to interpret the dynamic changes in gene expression and chromatin accessibility that occur in WT and *Mef2c* KO cells as they differentiate through each lineage.

### Cloning of Reporter Constructions for Zebrafish Transgenesis

To identify candidate enhancer regions to clone, we utilized our E8.5 ArchR dataset containing integrated chromatin accessibility and gene expression data. We identified all peaks that lost accessibility in the *Mef2c* KO (3,224), then filtered to the peaks that demonstrated altered accessibility in only one of the three heart tube segments (2,549), and that contained a MEF2 binding motif (1,059). Next, we utilized ArchR’s Peak2Gene functionality and filtered down to peaks correlated with DEGs in the *Mef2c* KOs (130) and were not located in promoter regions (119). Finally, from this list of 119 candidates, we selected 12 of the highest priority to screen based on the Log2FC value of the change in peak accessibility and linkage to genes of interest.

To generate the Tg(*MVEB:egfp*)^sfc^^26^ zebrafish lines, 650-900 nucleotide fragments containing the candidate enhancers were cloned by PCR from mouse genomic DNA.

Candidate nucleotide sequences and the primers used to amplify them are listed in Supplementary Table 6. The resulting fragments were cloned into the Tol2 transgenic vector E1b-eGFP-Tol2^30^ (RRID: Addgene_37845) using the In-Fusion Cloning system (Takara Bio). All plasmid sequences were confirmed by long-read sequencing before zebrafish injection (Plasmidsaurus). We were unable to successfully clone one of our twelve candidates (MVEB10) due to a highly repetitive “GT” sequence, so our final screen consisted of eleven candidate enhancers.

### Zebrafish Transgenesis Injections, Screening, and Imaging

All zebrafish experiments were reviewed and approved by the UCSF Institutional Animal Care and Use Committee and were performed in accordance with the Public Health Service Policy on the Humane Care and Use of Laboratory Animals. Zebrafish were raised under standard laboratory conditions at 28°C. The outbred wildtype zebrafish EKW (*Danio rerio-Ekwill*) and Tg(*cmlc2:mCherry*)^s890^ lines have been described previously^57^. One-cell zebrafish embryos obtained from Tg(*cmlc2:mCherry*)^s890^ and wildtype EKW (Ekwill strain) parents were injected with 1nl of injection mix containing 15pg of candidate-enhancer-eGFP and 15pg of ß-actin-BFP reporter plasmids along with 30pg of *Tol2* mRNA. The ß -actin promoter drives expression of BFP strongly in somite muscles and was used as a positive injection control. As a negative control, we also injected and screened embryos with an empty reporter construct containing only the minimal E1b promoter and eGFP. Injected embryos were screened at 24 and 72 hours post-fertilization (hpf) for eGFP and BFP expression. Representative embryos were imaged for eGFP expression as a readout of candidate enhancer activity and for mCherry expression as a myocardial marker. Enhancer activity was scored as a percentage of embryos expressing eGFP in the heart compared to the total number of BFP-positive embryos. A minimum of at least 50 BFP+ embryos were screened per construct, except for MVEB1, for which only 17 BFP+ embryos were obtained. The exact numbers screened for each construct are displayed in Extended Data Fig. 5b.

### Comparative Analyses with Published Datasets

NR2F2 and GATA4 target genes were downloaded from the Harmonizome^40^ and TFLink^41^ databases, respectively. Fisher’s exact test was used to determine the Odd’s Ratios and p-values for identifying NR2F2 and GATA4 targets among genes upregulated in *Mef2c* KO IFT-CMs at E8.5. Publicly available ChIP-seq datasets were downloaded from the Gene Expression Omnibus (GEO) in wig or bigwig format for MEF2C (GSE124008)^20^, H3K27ac (GSE52386)^27^, GATA4 (GSE52123)^39^, and NR2F2 (GSE46498)^38^. Datasets provided in wig format were converted to bigwig format using the Galaxy^58^ web tool wigToBigWig. CrossMap^59^ was used together with chain files from the UCSC Genome Browser^60^ to lift genome annotations from mm9 to mm10 reference genome assemblies, and vice versa. For the browser tracks in Fig. 5b-c, the MEF2C and H3K27ac ChIP-seq bigwigs were lifted from mm9 to mm10, the Galaxy^58^ web tool bigwigAverage was used to generate a single average MEF2C and H3K27ac occupancy track from the individual replicate tracks, and the results were displayed along with our pseudobulked snATAC-seq data using the IGV software package^55^. For the heat maps displayed in Extended Data Fig. 6b-c, we first created Browser Extensible Data (BED) format files for the gained DARs in *Mef2c* KO IFT-CMs containing NR or GATA motifs. We then lifted these BED regions from mm10 to mm9 and used the Galaxy^58^ web tools computeMatrix and plotHeatmap with the downloaded NR2F2 and GATA4 ChIP-seq data to visualize NR2F2 and GATA4 occupancy at the DARs. At the computeMatrix step, missing data was converted to zeroes. At the plotHeatmap step, kmeans clustering was used to generate two clusters of DARs (effectively “occupied” and “not occupied”) and these were sorted by decreasing mean value prior to being plotted.

### Gene Regulatory Network Inference and Analysis

We used CellOracle^43^ to infer GRNs for WT and *Mef2c* KO IFT-CMs at E8.5 and E9. The GRN inference process included the following steps:

### Base GRN Construction

We used OFT-CM, V-CM, IFT-CM, and A-CM snATAC-seq data to construct base GRNs that define the accessible TF binding motifs near genes for each of the following conditions: E8.5-WT, E8.5-KO, E9-WT, and E9-KO. Pooling the OFT-CM, V-CM, and A-CM/IFT-CMs at each timepoint mitigated artifacts caused by data sparsity. The binarized PeakMatrix output from ArchR was used as an input to Cicero’s^61^ run_cicero() function (with default parameters) to compute co-accessibility scores among peak pairs within 500 kb. Output scores were from -1 to 1, where higher scores indicate stronger co-accessibility. Each dataset generated over 20 million peak pairs for co-accessibility comparisons. We annotated transcription start sites (TSSs) with their corresponding genes for accessible peaks using the motif_analysis.get_tss_info() function in CellOracle. Unlabeled peaks were considered potential regulatory elements. Subsequently, we applied motif_analysis.integrate_tss_peak_with_cicero() to link unlabeled peaks with genes, filtering for peak-to-gene pairs with co-accessibility scores > 0.8. TF binding motifs within each 500 bp peak were identified using the motif_analysis.TFinfo() function in CellOracle, which converts peak IDs into DNA sequences using the mm10 reference genome. TF motif scanning used the gimme.vertebrate.v5.0^62^ library to produce a set of TF motifs present in accessible regions within 500kb of each gene. Finally, differentially accessible chromatin regions in KO-vs-WT were determined for each timepoint (FDR < 0.01) and inaccessible regions were pruned from the base GRNs. The TFs that bind to motifs present in regions that are co-accessible with and within 500 kb of the TSS of each gene are inferred to have the potential to regulate that gene.

### GRN Inference

Using CellOracle, we constructed GRNs in which edge strengths are quantified based on the TF-to-gene correlation, constrained by the accessible regions defined in the base GRN. We inferred GRNs for E8.5 WT, E8.5 KO, E9 WT, and E9 KO IFT-CMs, using the computed base GRNs to determine permissible TF-to-gene links. GRN inference used the top 5,000 differentially expressed genes in the snRNA-seq dataset. We used the CellOracle oracle.get_links() function, with alpha set to 1,000 and otherwise default parameters, fitting linear models with L2 weight regularization to derive coefficients relating TF expression to target gene expression. The Bagging Ridge model was used to fit coefficients, providing posterior distributions for each TF-to-gene relationship. Positive coefficients indicated activating, while negative coefficients indicated inhibiting relationships. We then filtered the inferred network to include 12,000 edges with the largest magnitude coefficients.

### Simulation of Mef2c Knockout Effects

To predict the effect of *Mef2c* KO on gene expression, we identified genes directly regulated by MEF2C in the E8.5 WT IFT-CM network. We predicted that reduced expression of *Mef2c* in the network would downregulate genes directly activated by MEF2C and upregulate genes directly inhibited by MEF2C in the E8.5 WT IFT-CM network. The simulation results were compared against differentially expressed genes in the E9 snRNA-seq IFT-CM experimental data (KO-vs-WT). Simulations were performed and prediction accuracy tested with varying Log2FC cutoffs for differentially expressed genes, as summarized in Supplementary Table 8.

### Network Visualization

Whole-network visualizations were generated using Gephi^63^, applying the Yifan Hu layout, with node sizes scaled by betweenness centrality. PyVis^64^ was used to visualize subnetworks that included manually selected TFs (*Mef2a, Mef2c, Nr2f2, Tbx5, Gata4, Meis2, Plagl1, Isl1, Pitx2, Tbx20, Hand2, Nkx2-5*) and the top 100 differentially expressed genes (ranked in ascending order by p value) identified by comparing the E8.5 WT and *Mef2c* KO IFT-CM snRNA-seq experimental data. Visualizations for direct interactions between TFs (*Nr2f2* and *Gata4*) and targets in the E8.5 WT and *Mef2c* KO IFT-CMs were developed using Python’s NetworkX package^65^, with a circular layout and edge widths scaled by coefficient values.

### Phenotypic Analysis of *Mef2c*^-/-^ and *Mef2c*^-/-^; *Nr2f2*^+/-^ Embryos

We conducted comparative phenotypic analysis of *Mef2c*^-/-^ and *Mef2c*^-/-^; *Nr2f2*^+/-^ embryos by performing immunofluorescent staining of cardiac Troponin T and analyzing the apparent heart tube phenotypes, initially unblinded to embryo genotype. We determined that the heart tube phenotypes ranged in apparent severity and fell into roughly three categories, with Category 1 being the most severely malformed and Category 3 the least severe. We then blinded ourselves to the embryo genotypes and re-examined the embryos, placing each into the appropriate category of severity.

## Supporting information

Supplementary Table Descriptions

## ACKNOWLEDGEMENTS

J.M.M-V. is grateful for assistance from Dr. Kavitha Rao in learning Seurat, as well as from Drs Ryan Corces and Christina Theodoris for assistance with ArchR and MIRA, respectively. J.M.M-V. is also grateful to all his colleagues in the Bruneau Lab for many insightful discussions, important feedback on directions of the project, and help with experiments. We thank the Gladstone Genomics Core for use and maintenance of the Agilent Bioanalyzer and 10x Genomics Chromium Controller. Portions of this work were performed on the Wynton HPC Co-Op cluster, which is supported by UCSF research faculty and UCSF institutional funds. The authors wish to thank the UCSF Wynton team for their ongoing technical support of the Wynton environment.

## AUTHOR CONTRIBUTIONS

J.M.M-V. and B.G.B. conceived and designed the experiments presented in this study. J.M.M-V. performed embryo dissections, whole-mount embryo immunostaining, whole-mount embryo FISH via HCR, embryo imaging, preparation of sequencing libraries, and data analysis via Seurat, ArchR, and MIRA. E.F.B. performed mouse colony maintenance and genotyping. J.M.M-V. and E.F.B. performed cloning of reporter constructs for zebrafish transgenesis. T.S. performed zebrafish transgenesis injections and imaging. A.P.C. performed GRN analyses using CellOracle and newly written code. B.G.B., B.L.B., and J.J.S. supervised and advised. J.M.M-V. prepared the figures and wrote the manuscript with input from the co-authors.

## DATA AND CODE AVAILABILITY

Raw and processed data for snRNA-seq and snATAC-seq datasets has been deposited in the NCBI Gene Expression Omnibus (GEO) under accession number GSE280587. Code used for multiomics analyses, GRNs, and plot generation is available in the GitHub repository associated with this manuscript: https://github.com/jmmuncie/Muncie-Vasic_2025.

