## Extended Data Figures for "MEF2C controls segment-specific gene regulatory networks that direct heart tube morphogenesis"

**Extended Data Figure 1**

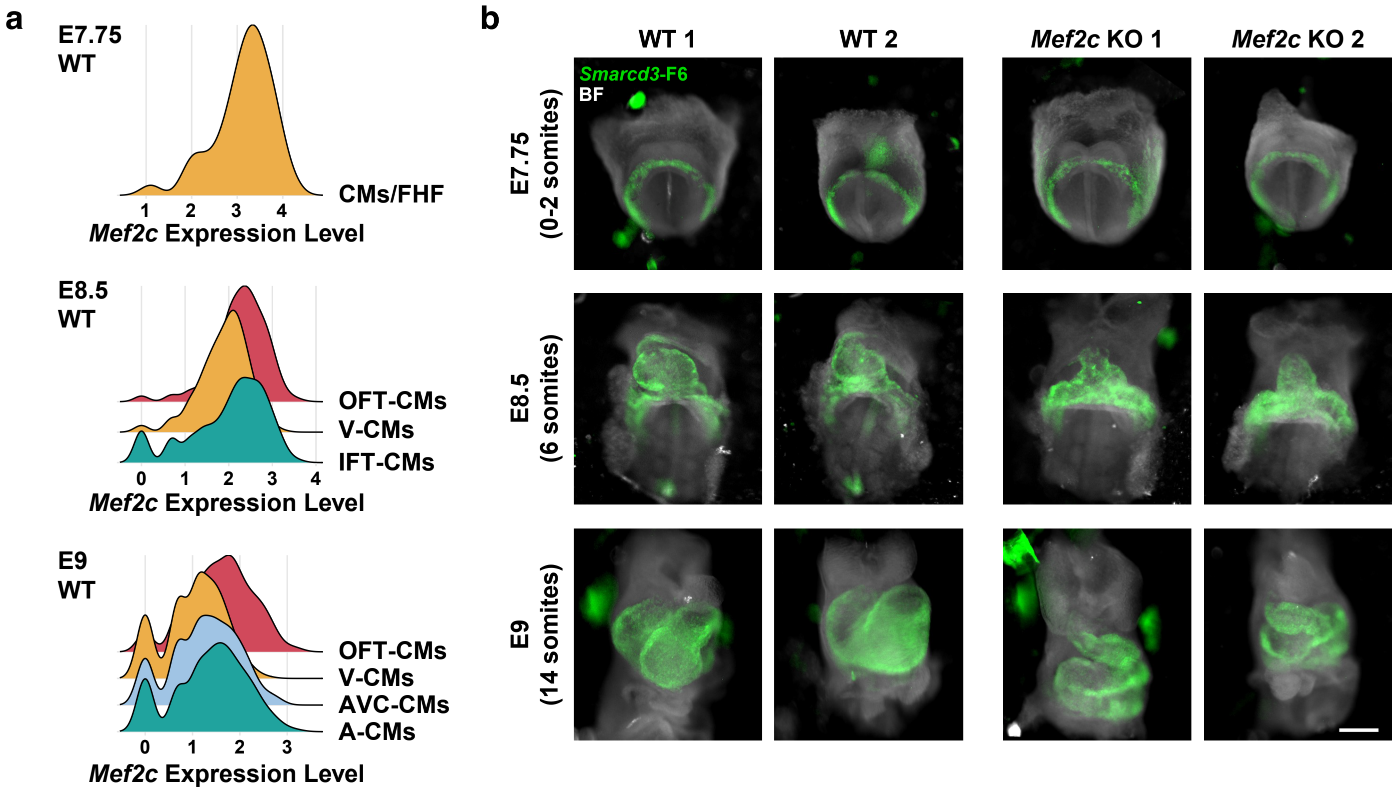


**Extended Data Figure 1:** a) Ridge plots displaying *Mef2c* expression in cardiac progenitors and CM subtypes in E7.75, E8.5, and E9 embryos. b) Images of all embryos collected for 10x Multiome (snRNA-seq and snATAC-seq) experiment. Cardiac progenitors are marked by the *Smarcd3*-F6-eGFP reporter transgene (green). BF, brightfield. Scale bar = 200 µm. CMs, cardiomyocytes; FHF, first heart field; OFT-CMs, outflow tract cardiomyocytes; V-CMs, ventricular cardiomyocytes; IFT-CMs, inflow tract cardiomyocytes; AVC-CMs, atrioventricular canal cardiomyocytes; A-CMs, atrial cardiomyocytes.

### Extended Data Figure 2

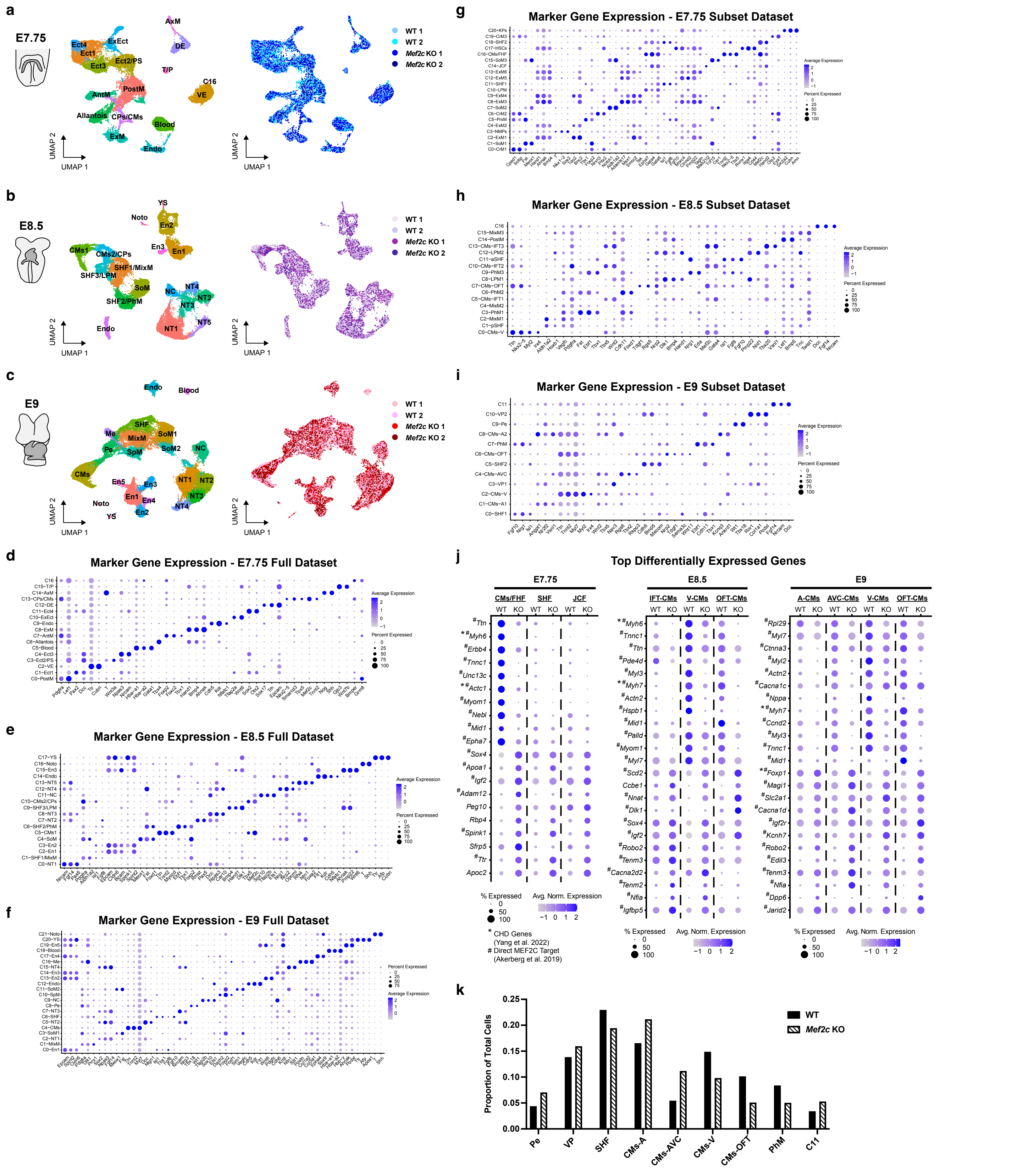


**Extended Data Figure 2:** a-c) UMAPs of full snRNA-seq datasets from E7.75 (a), E8.5 (b), and E9 (c) embryos labeled by cell type (left) and genotype/sample ID (right). d-f) Dot plots displaying expression of key marker genes used to identify and label cell types in the full snRNA-seq datasets for E7.75 (d),

E8.5 (e), and E9 (f) embryos. g-i) Dot plots displaying expression of key marker genes used to identify and label cell types in the subset datasets consisting of cardiac and related mesoderm cells from E7.75 (g), E8.5 (h), and E9 (i) embryos. j) Dot plots displaying gene expression of the top DEGs at E7.75, E8.5, and E9. k) Bar plot quantifying the proportion of total cells per cell type for WT and *Mef2c* KO cells in the E9 subset dataset. Ect, ectoderm; ExEct, extraembryonic ectoderm; PS, primitive streak; AntM, anterior mesoderm; PostM, posterior mesoderm; CPs, cardiac progenitors; CMs, cardiomyocytes; ExM, extraembryonic mesoderm; AxM, axial mesoderm; DE, definitive endoderm; T/P, trophoblast and placental progenitors; VE, visceral endoderm; Endo, endothelium; SHF, second heart field; MixM, mixed mesoderm; LPM, lateral plate mesoderm; SoM, somitic mesoderm; PhM, pharyngeal mesoderm; YS, yolk sac; Noto, notochord; En, endoderm; NT, neural tube; NC, neural crest; SpM, splanchnic mesoderm; Me, mesenchyme; Pe, proepicardium.

#### Extended Data Figure 3

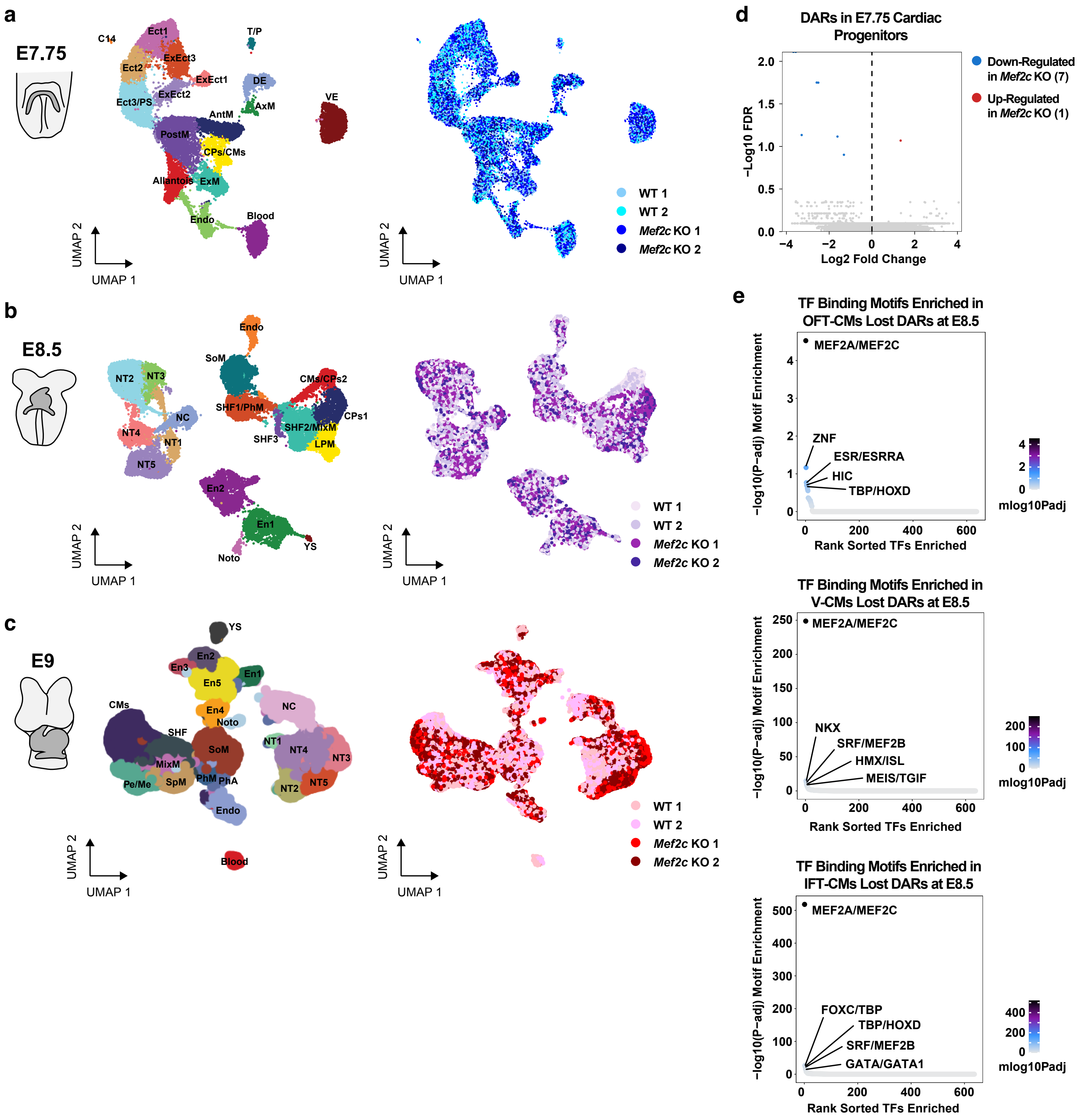


**Extended Data Figure 3:** a-c) UMAPs of full integrated snRNA-seq and snATAC-seq datasets from E7.75 (a), E8.5 (b), and E9 (c) embryos labeled by cell type (left) and genotype/sample ID (right). d) Volcano plot displaying results of differential accessibility testing (*Mef2c* KO relative to WT) in cardiac progenitors at E7.75. e) TF binding motif enrichment analysis for lost DARs (*Mef2c* KO relative to WT) in OFT-CMs (top), V-CMs (middle), and IFT-CMs (bottom) at E8.5. Ect, ectoderm; ExEct, extraembryonic ectoderm; PS, primitive streak; AntM, anterior mesoderm; PostM, posterior mesoderm; CPs, cardiac progenitors; CMs, cardiomyocytes; ExM, extraembryonic mesoderm; AxM, axial mesoderm; DE, definitive endoderm; T/P, trophoblast and placental progenitors; VE, visceral endoderm; Endo, endothelium; SHF, second heart field; MixM, mixed mesoderm; LPM, lateral plate mesoderm; SoM, somitic mesoderm; PhM, pharyngeal mesoderm; PhA, pharyngeal arches; YS, yolk sac; Noto, notochord; En, endoderm; NT, neural tube; NC, neural crest; SpM, splanchnic mesoderm; Me, mesenchyme; Pe, proepicardium.

#### Extended Data Figure 4

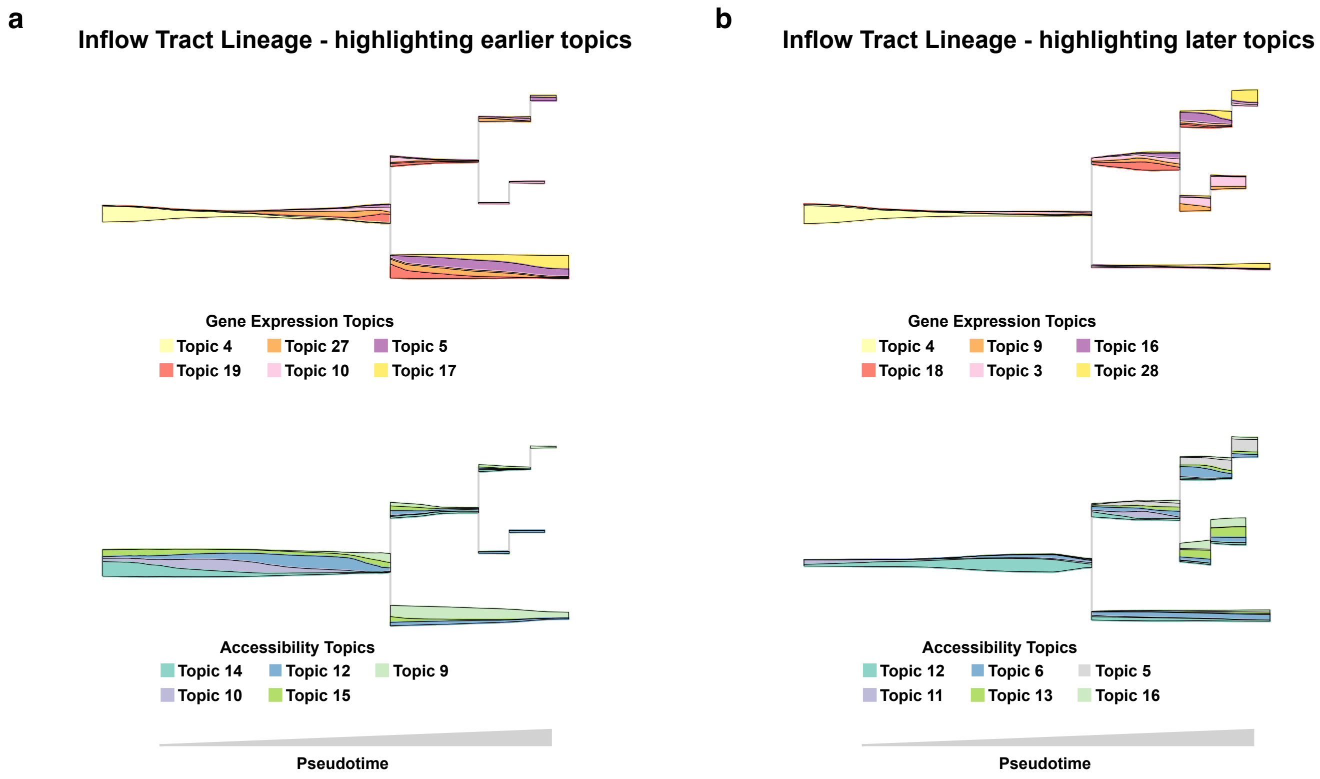


**Extended Data Figure 4:** a-b) Stream plots displaying the flow of gene expression (top) and chromatin accessibility (bottom) topics in the inflow tract lineage cells, as in Fig. 4m, with additional topics displayed to highlight differences in earlier (a) and later (b) branch points along the trajectory.

**Extended Data Figure 5**

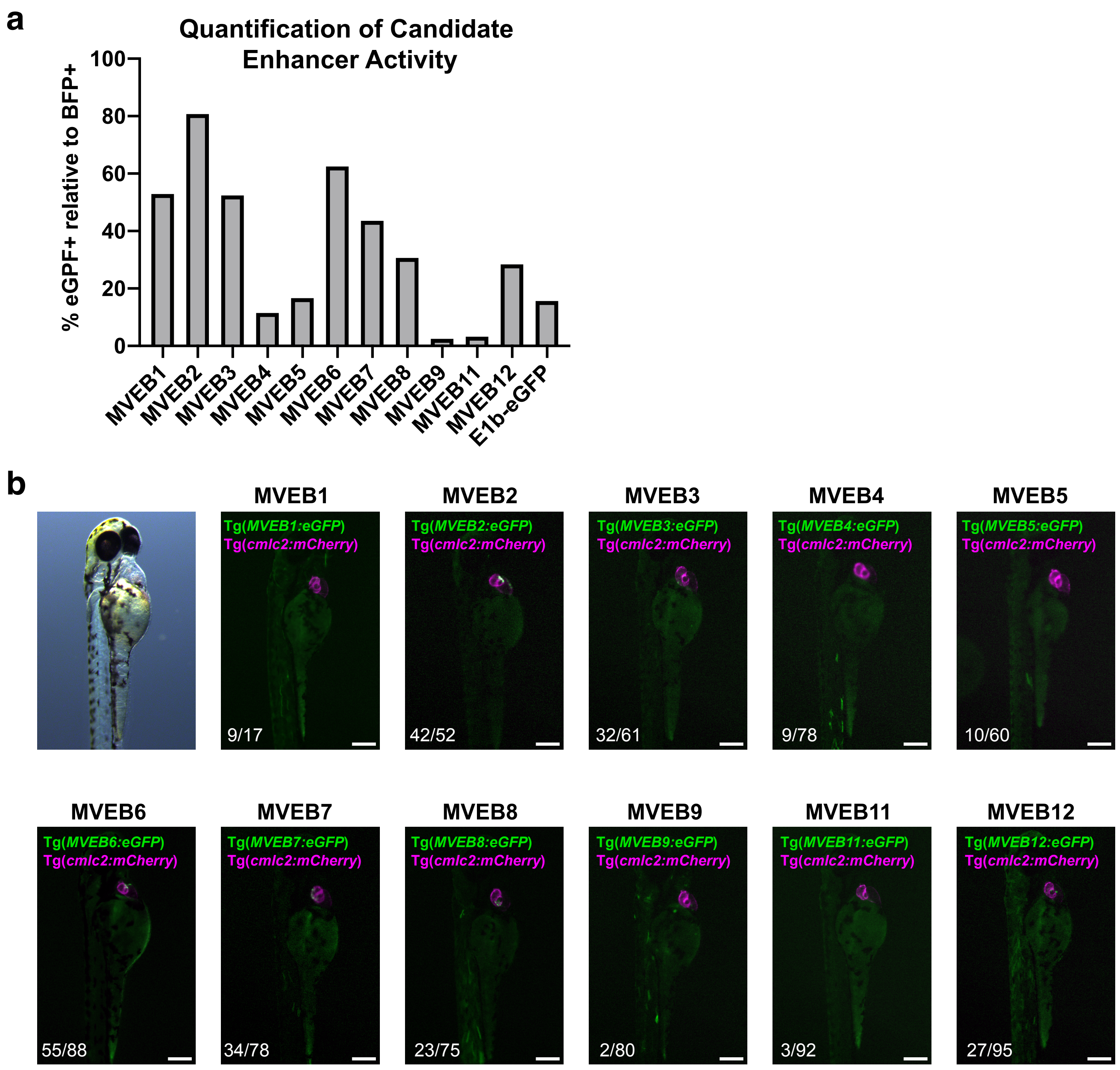


**Extended Data Figure 5:** a) Quantification of the percentage of BFP+ zebrafish embryos that exhibited eGFP expression in the heart at 72 hpf. b) Representative lateral views of zebrafish embryos at 72 hpf injected with candidate enhancers. Representative brightfield image (left) demonstrates the orientation and scale of the fluorescent images. Numbers in the bottom-left corner indicate the number of embryos with eGFP expression in the heart relative to the number of BFP+ embryos. Scale bars = 200 μm.

#### Extended Data Figure 6

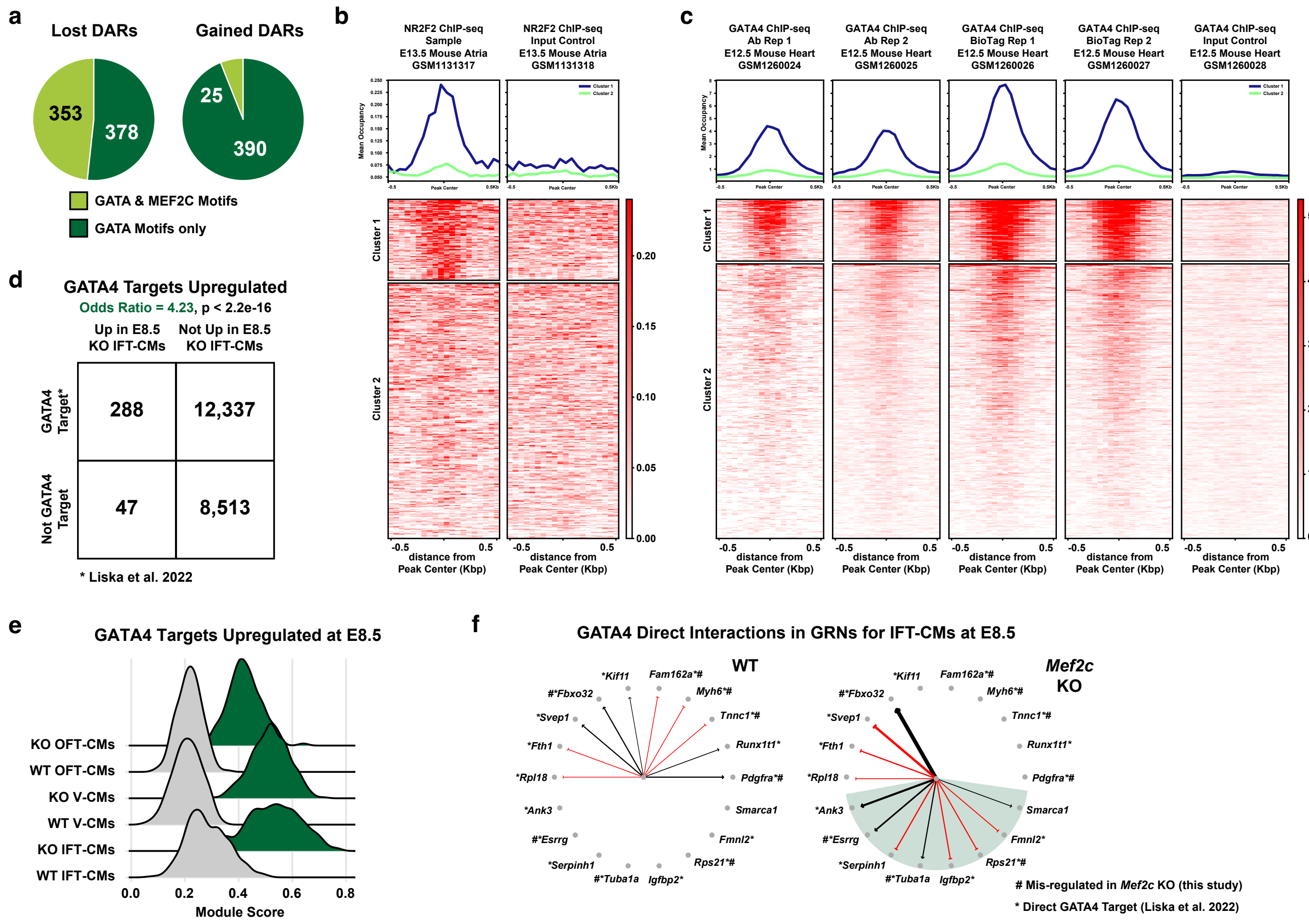


**Extended Data Figure 6:** a) Pie charts showing the proportion of gained or lost DARs (*Mef2c* KO relative to WT) containing GATA and MEF2C motifs or only GATA motifs. b-c) Heatmaps and average occupancy profiles of NR2F2 (b) and GATA4 (c) ChIP-seq data[38,39] at gained DARs (*Mef2c* KO relative to WT) in E8.5 IFT-CMs that contained NR (b) or GATA (c) motifs. kmeans clustering was used to generate two clusters of DARs representing occupied (cluster 1) and non-occupied (cluster 2) regions. d) Odds ratio analysis for GATA4 target genes[41] amongst DEGs up-regulated in *Mef2c* KO IFT-CMs. p-value calculated using Fisher's exact test. e) Ridge plot displaying module scores for up-regulated GATA4 targets in *Mef2c* KO and WT heart tube segments. f) Visualization of direct GATA4 interactions in the WT and *Mef2c* KO E8.5 IFT-CM GRNs. Direct interactions that occur upon *Mef2c* KO are highlighted. #, Mis-regulated DEG in *Mef2c* KO IFT-CMs at E8.5; *, Direct target of GATA4[41].

#### Extended Data Figure 7

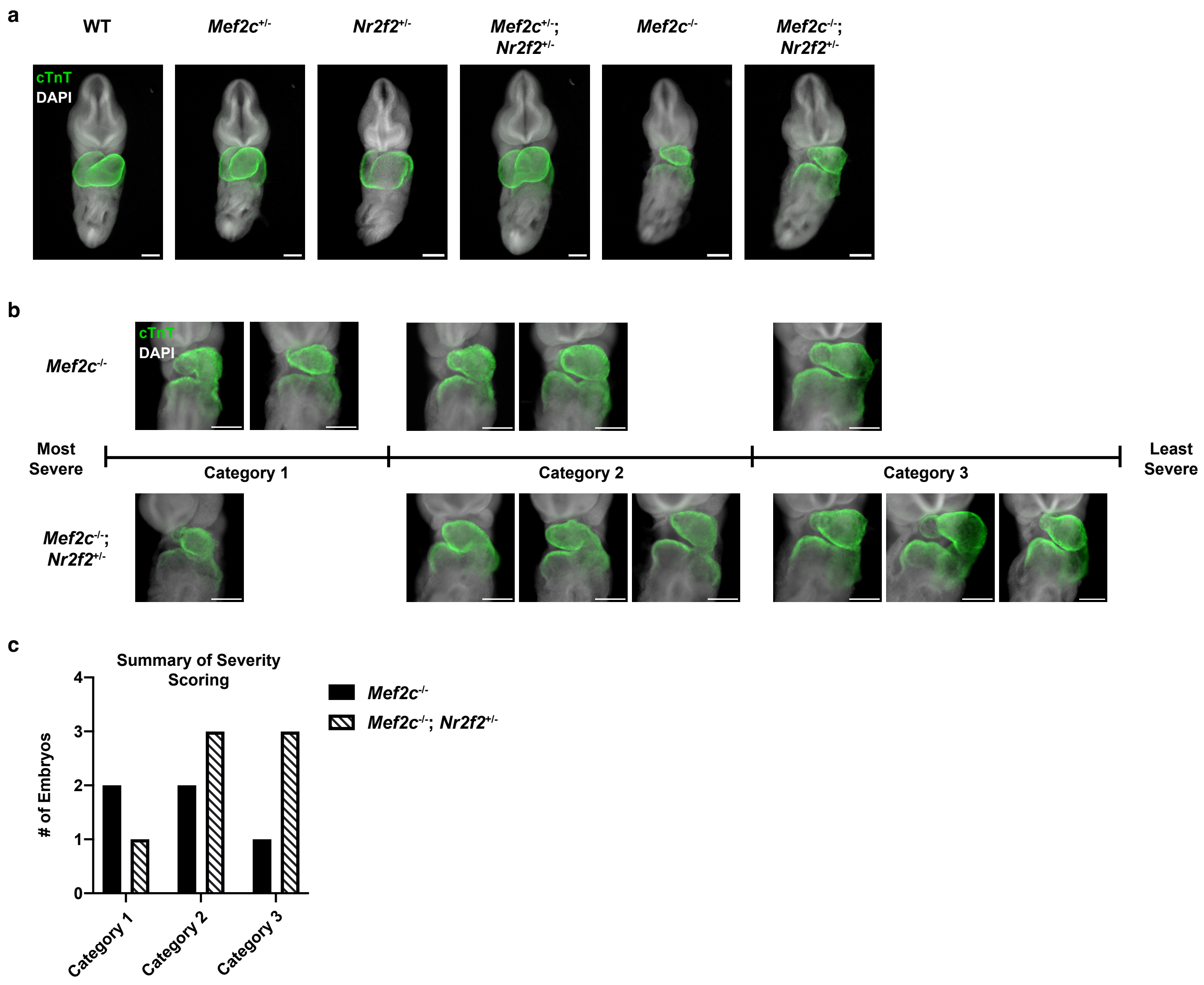


**Extended Data Figure 7:** a) Immunofluorescent staining of cardiac Troponin T (cTnT, green) in representative E9.5 (18-24 somites) embryos from each genotype collected from *Mef2c*$^{+/-}$;*Nr2f2*$^{+/-}$ to *Mef2c*$^{+/-}$ crosses. b) Immunofluorescent staining of cardiac Troponin T (cTnT, green) in all *Mef2c*$^{-/-}$ (n=5) and *Mef2c*$^{-/-}$;*Nr2f2*$^{+/-}$ (n=7) embryos collected at E9.5 (18-24 somites) from 7 independent litters, arranged by severity of heart tube phenotype. Scale bars = 200 μm. c) Bar plot summarizing the heart tube phenotype severity scoring of *Mef2c*$^{-/-}$ and *Mef2c*$^{-/-}$;*Nr2f2*$^{+/-}$ embryos shown in (b).
