## Supplementary Table Descriptions for "MEF2C controls segment-specific gene regulatory networks that direct heart tube morphogenesis"

**Supplementary Table 1:** Marker genes used to identify and label cell types in full snRNA-seq datasets for E7.75, E8.5 and E9 embryos.

**Supplementary Table 2:** Marker genes used to identify and label cell types in subset snRNA-seq datasets consisting of cardiac progenitors and related mesoderm in E7.75, E8.5 and E9 embryos.

**Supplementary Table 3:** Differential gene expression results (*Mef2c* KO-vs-WT) for cardiac progenitors and cardiomyocyte subtypes in E7.75, E8.5, and E9 embryos.

**Supplementary Table 4:** Summary of Peak2Gene, DAR, and DEG information for all Peak2Gene links in E8.5 OFT-CMs, V-CMs, and IFT-CMs.

**Supplementary Table 5:** MIRA topic compositions consisting of the top 200 genes for each gene expression topic and motif enrichment scores for the chromatin accessibility topics.

**Supplementary Table 6:** List of high-priority candidate enhancers for zebrafish transgenesis screen.

**Supplementary Table 7:** NR2F2 and GATA4 target genes that were upregulated in E8.5 *Mef2c* KO IFT-CMs.

**Supplementary Table 8:** Summarized prediction accuracy of simulated *Mef2c* knockdown in CellOracle GRNs with varied Log2FC cutoffs for differentially expressed genes.
